# High vitamin B_2_ production by the *Lactiplantibacillus plantarum* B2 strain carrying a mutation in the aptamer P1 helix of the FMN-riboswitch that regulates expression of the bacterial *rib* operon

**DOI:** 10.1101/2021.05.20.444923

**Authors:** Inés Ripa, José Ángel Ruiz-Masó, Nicola De Simone, Pasquale Russo, Giuseppe Spano, Gloria del Solar

## Abstract

Manufacturing of probiotics and functional foods using lactic acid bacteria that overproduce vitamin B_2_ has gained growing interest due to ariboflavinosis problems affecting populations of both developing and affluent countries. Two isogenic *Lactiplantibacillus plantarum* strains, namely a riboflavin-producing parental strain (UFG9) and a roseoflavin-resistant strain (B2) that carries a mutation in the FMN-aptamer of the potential *rib* operon riboswitch, were analyzed for production and intra- and extracellular accumulation of flavins, as well as for regulation of the *rib* operon expression. Strain B2 accumulated in the medium one of the highest levels of riboflavin+FMN ever reported for lactic acid bacteria, exceeding by ~25 times those accumulated by UFG9. Inside the cells, concentration of FAD was similar in both strains, while that of riboflavin+FMN was ~6-fold higher in B2. Mutation B2 could decrease the stability of the aptamer’s regulatory P1 helix even in the presence of the effector, thus promoting the antiterminator structure of the riboswitch ON state. Although the B2-mutant riboswitch showed an impaired regulatory activity, it retained partial functionality being still sensitive to the effector. The extraordinary capacity of strain B2 to produce riboflavine, together with its metabolic versatility and probiotic properties, can be exploited for manufacturing multifunctional foods.

## 1 Introduction

Riboflavin (RF, also named vitamin B_2_) is the precursor of the coenzymes flavin mononucleotide (FMN) and flavin adenine dinucleotide (FAD) (Fig. S1B) that are involved in multiple redox reactions implied in the electron transport respiratory chain, in the metabolism of carbohydrates, lipids, proteins and some toxin and drugs, as well as in the oxidative stress response and in the activation of other B-group vitamins. Apart from redox reactions, flavoproteins also participate in processes such as light sensing, body development, DNA repair and circadian cycling (García-Angulo, 2017). Humans cannot synthesize RF, and thus it must be supplied from the diet and/or the gut microbiota (LeBlanc et al., 2013). Although the average daily intake of RF within developed countries is in general above the recommended dietary allowance, a suboptimal RF status is a widespread problem among certain groups particularly susceptible to this condition (e.g. teenagers, elderly people, pregnant women, alcohol abusers, people with long-standing infections or liver disease; Fabian et al., 2012; Mensink et al., 2013; Mazur-Bialy et al., 2015). In underdeveloped countries, there is a higher prevalence of ariboflavinosis due to widespread undernutrition, especially in children (Sherwood, 2018).

In contrast to humans and other animals, plants, fungi and most prokaryotes are able to synthesize riboflavin *de novo* from GTP and ribulose 5-phosphate. All organisms, however, are able to convert RF into FMN and FAD (Fig. S1B). Bacteria highly differ in the presence, genetic organization and regulation of the genes involved in the RF biosynthetic pathway and in the transport of flavins, even among phylogenetically related species. Most bacteria carry genes involved in the regulated biosynthesis of RF, whereas others are RF auxotrophs. These latter necessarily rely on specialized flavin transport systems for the uptake of exogenous RF. Specific importer systems are also present in some RF-prototrophic bacteria, which are hence enabled to avoid the metabolic cost of synthesizing riboflavin when it is present in the environment (Abbas and Sibirny, 2011; García-Angulo, 2017).

In Gram-positive bacteria, expression of the *rib* operon (Fig. S1A) is regulated by FMN riboswitch-mediated transcriptional attenuation (Thakur et al., 2016). These FMN riboswitches are RNA elements located at the 5’ untranslated region of the *rib* operon mRNA that consist of a conserved FMN-sensing aptamer domain (the so-called RFN element) and an expression platform (or regulatory domain) exhibiting alternative terminator (OFF state) and antiterminator (ON state) conformations (Abbas and Sibirny, 2011). Binding of FMN to the RFN element in the nascent mRNA results in a shift in the conformation of the expression platform towards the OFF state of the riboswitch. The effect of FMN binding to the aptamer on the conformation of the downstream expression platform was studied by analyzing the fraction of full-length and terminated products of *in vitro* transcription of the untranslated leader region of the *Bacillus subtilis rib* operon (Winkler et al., 2002; Wickiser et al., 2005). Among lactic acid bacteria (LAB), the most detailed genetic and functional study of the *rib* operon riboswitch has been performed in *Lactococcus lactis* (Burgess et al., 2004), where the transcription start site of the *rib* operon and the DNA region encoding the leader mRNA were determined, and the RFN element and terminator, antiterminator and anti-antiterminator sequences identified (Fig. S1A). Also, the *in vivo* regulatory activity of this riboswitch was studied by cloning the DNA that encodes the mRNA leader region of the *rib* operon in a promoter-probe vector and analyzing the expression of the β-galactosidase reporter gene in lactococcal cells grown in chemically-defined medium either supplemented or not with RF or FMN (Burgess et al., 2004). The coexistence of RF uptake systems and RF biosynthetic pathways makes many Gram-positive bacteria naturally sensitive to roseoflavin, an analogue of riboflavin produced by *Streptomyces davaonensis* and *Streptomyces cinnabarinus* that is imported into the target cell, where it can bind to the aptamer of the FMN riboswitch promoting its terminator conformation and hence the shutting down of the *rib* operon. Exposition of RF-producing microorganisms to the selective pressure of roseoflavin has been reported as a strategy to select LAB with RF-overproducing phenotype (Burgess et al., 2004; Abbas and Sibirny, 2011). In particular, by using this approach several derivative LAB strains able to overproduce RF to different extents have been selected that belong to *L. lactis, L. plantarum, Limosilactobacillus fermentum*, and *Leuconostoc mesenteroides* species (Burgess et al., 2004, 2006; Capozzi et al., 2011; Russo et al., 2014a; Mohedano et al., 2019). Single-base mutations and deletions affecting several regions of the RFN element, as well as deletions and insertions within the expression platform have been reported in these RF-overproducing strains (Burgess et al., 2004, 2006; Russo et al., 2014a; Ge et al., 2020). In spite of the phenotype of vitamin B2 overproduction, impaired regulatory functionality of the *rib* operon riboswitch has only been directly demonstrated for one lactococcal mutant strain (Burgess et al., 2004).

For several decades and until recently, commercial RF was mainly produced by chemical or chemoenzymatic processes, which involved use of toxic compounds and high costs of waste disposal. Hence, environmental and economic considerations led to the replacement of organic synthesis of RF by its biotechnological production (Thakur et al., 2016; Revuelta et al., 2017). Currently, vitamin B_2_ is exclusively produced by economical and eco-efficient biotechnological processes based on RF-overproducing strains of the bacterium *B. subtilis* and the filamentous fungus *Ashbya gossypii*, which yield >20 g/L of riboflavin (Revuelta et al., 2017; Acevedo-Rocha et al., 2019). Recently, there has also been a growing use of LAB able to synthesize B-group vitamins (especially RF) for *in situ* food bio-fortification, which reduces the production costs and increases the vitamin bioavailability (Thakur et al., 2016). Compared to *B. subtilis* and *A. gossypii*, the RF yield in overproducing LAB strains is quite low. However, the metabolic versatility and adaptability to fermentation processes of this group of bacteria make them most suitable for the elaboration of vitamin B2-enriched fermented or processed food. *Lactiplantibacillus plantarum* (formerly known as *Lactobacillus plantarum* following an updated taxonomical reclassification of LAB; Zheng et al., 2020), is a food-grade bacterium widely employed in the food industry as starter and/or as probiotic. Previously, *L. plantarum* B2 (*Lp* B2), a roseoflavin-resistant mutant overproducing vitamin B2 that exhibited one of the highest RF production among LAB, was characterized for its probiotic potential *in vitro* (Arena et al., 2014), as well as in a zebrafish *in vivo* model (Russo et al., 2015a). These assessments boosted application studies of *Lp* B2 in the food industry, showing its capacity to produce high levels of vitamin B_2_ during fermentation of oat-based foods (Russo et al., 2016) and soymilk (Zhu et al., 2020), as well as its adequacy for the production of probiotic fresh-cut pineapple (Russo et al., 2014b) and cantaloupe (Russo et al., 2015b).

In this work, we have studied the capacities of the isogenic wild-type (WT) UFG9 (*Lp* UFG9) and mutant *Lp* B2 strains of *L. plantarum* to produce flavin compounds and to regulate this production. The real-time accumulation of flavins in the culture medium during the growth of strains UFG9 and B2 was analyzed. In addition, a method was implemented that allowed the differential quantification of the intracellular and extracellular amounts of both RF+FMN and FAD. Moreover, an *in silico* analysis of the 5’ untranslated region of the *rib* operon of *Lp* UFG9 allowed us to identify the main sequence and structural elements potentially involved in the shift between the ON and OFF conformational states of the riboswitch that regulates expression of the genes encoding the RF biosynthetic pathway. Finally, the regulatory activity of the WT and B2-mutant riboswitches was analyzed *in vivo* by cloning the corresponding DNA fragments into promoter-probe vector pRCR, which carries the mCherry-encoding reporter gene, and quantifying the fluorescence emitted when the *Lp* UFG9 and *Lp* B2 host cells were grown in the presence or in the absence of RF.

## 2 Results

### 2.1 Mutant strain *Lp* B2 renders much higher amounts of flavins than its isogenic WT strain

Production of flavin compounds by strains *Lp* B2 and *Lp* UFG9 was detected in real time by monitoring the fluorescence (λ_ex/em_ 450/530 nm) of the cultures during the bacterial growth in medium CDM, which is directly proportional to the flavin concentration (Fig. 1A and B; Mohedano et al., 2019). In this approach, RF was used as the standard (Fig. 1B), since this is the flavin expected to accumulate in roseoflavin-resistant mutants (Burgess et al., 2006). Thus, the values of fluorescence intensity emitted by the cultures were assigned with the corresponding RF concentrations (Fig. 1A). Both *L. plantarum* isogenic strains grew similarly, but only B2 showed significant production of flavins, which accumulated in the cultures of the mutant strain and reached a plateau at the stationary phase (Fig. 1A). Maximal yield of flavins in the *Lp* B2 cultures was equivalent to ~3.7 mg/L RF, a concentration more than 20-fold higher than that detected in the cultures of the UFG9 WT strain (equivalent to ~0.16 mg/L RF; Fig. 1).

**Fig. 1.**
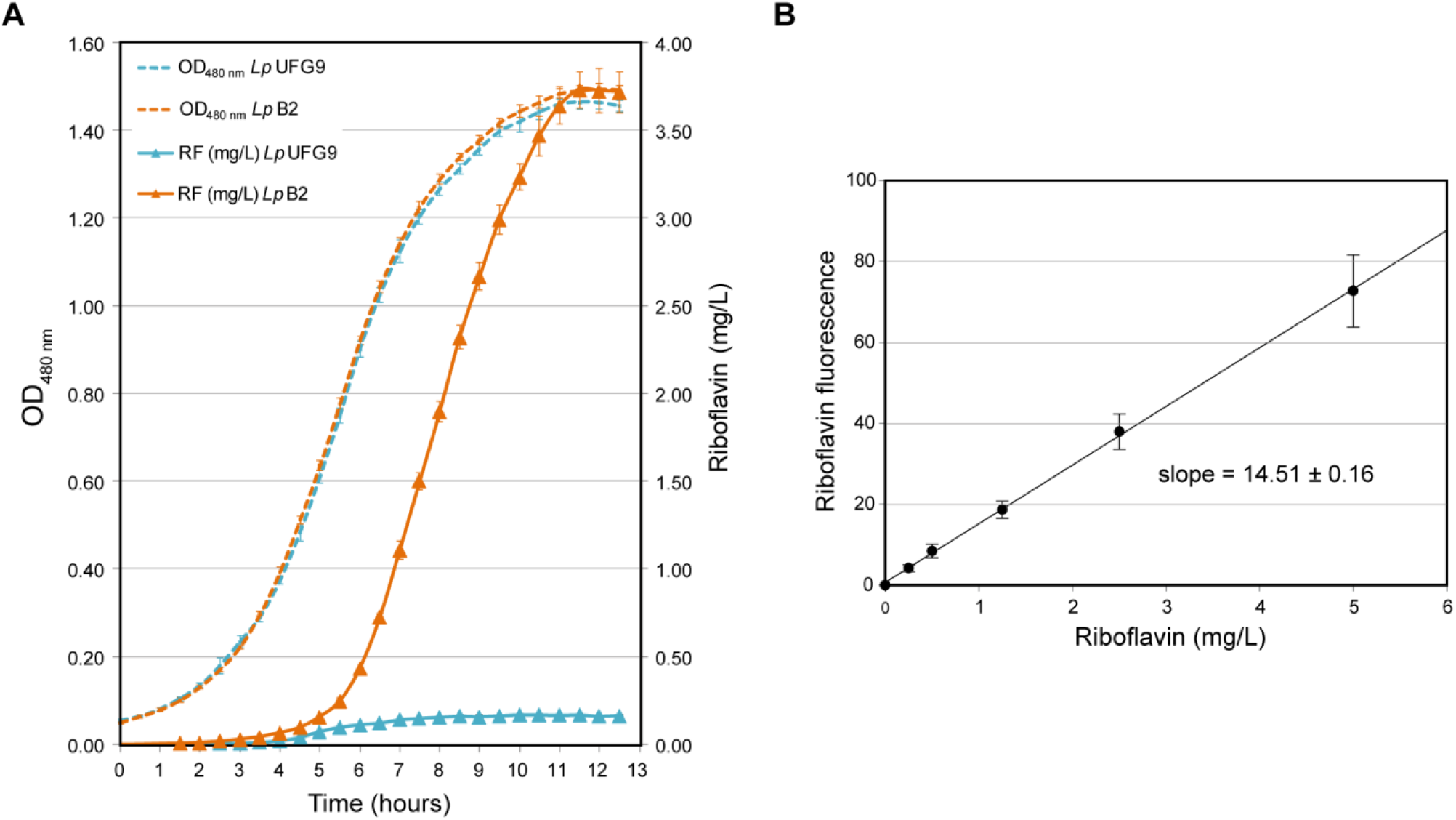
Real time monitorization of riboflavin production. **A)** Comparative analysis of the growth curves of *Lp* UFG9 and *Lp* B2 in CDM, monitored at an OD of 480 nm, and the concentration of RF equivalents in the same cultures, as calculated from the calibration curve depicted in panel B. Plots display representative growth and RF concentration curves of each strain obtained from one of at least three independent experiments (biological samples), with the symbols and vertical bars representing the average and errors of three technical triplicates. **B)** Calibration curve made by using 6 standard solutions with different RF concentrations, which did result in a highly linear relationship (R^2^ of 0.99). The experimental data were fitted to a linear regression model of slope = 14.51 ± 0.16 (intrinsic fluorescence of the RF). The calibration curve was elaborated from three independent experiments.

We next investigated whether the real-time detected flavins were located inside or outside the cells in cultures of *Lp* B2 and *Lp* UFG9. In the WT strain, similar fluorescence intensity values were observed in the supernatant and in the washed cells. In contrast, the supernatant of the *Lp* B2 culture displayed a fluorescence intensity ~6.5-fold higher than the corresponding washed cells. Moreover, the fluorescence intensities corresponding to the washed cells and to the supernatant of the *Lp* B2 culture were about 5- and 33-fold higher, respectively, than those of the *Lp* UFG9 culture (Table S3). These results show that: i) this method enables the real-time joint detection of intra- and extracellular flavins, and ii) a main fraction of the flavin excess in the *Lp* B2 mutant strain is exported to the medium. Curiously, the washed cell suspension and the supernatant did not add the fluorescence intensity of the corresponding uncentrifuged culture, which was ~1.4-1.5-fold higher than that sum for either strain (Table S3). This apparent inconsistency may arise from a positive interference of the cells that potentiates the fluorescence of the flavins present in the culture supernatant, or from the undesired loss of flavins weakly bound to the cell surface during the pellet washing step. To distinguish between these alternative explanations, the cellular pellets washed with PBS were resuspended in their own supernatants. The fluorescence intensities of the reconstituted cultures became similar to those of the initial cultures, pointing to a positive interference of the cells, beyond the fluorescence due to their flavin content, on the fluorescence intensity of the supernatants. Thus, this real-time method, although underestimates FAD due to its lower intrinsic fluorescence compared with RF and FMN, seems to entail interference artifacts by the cells that result in overestimation of the flavin concentration.

### 2.2 *Lp* B2 secretes much greater amounts of RF+FMN and FAD and accumulates a higher intracellular concentration of RF+FMN than its isogenic *Lp* UFG9 WT strain

In order to differentially quantify the distinct flavin compounds located within the cells or secreted to the medium by *Lp* UFG9 and *Lp* B2, we set up a fluorimetric method based on the lower intrinsic fluorescence of FAD relative to RF or FMN at neutral pH and on the acid- and temperature-dependent conversion of FAD into FMN and RF. Hence, this method allows distinguishing between RF+FMN and FAD by analyzing the increase in fluorescence of samples that have been subjected to the acidic treatment. A similar approach was previously used in *L. lactis* to analyze the intracellular content of FAD, which was expressed as the amount of RF with the same fluorescence (Chen et al., 2013). In contrast, we obtained the standard curves correlating the fluorescence intensity with the concentration of either RF or FAD (Fig. S2) and used a modified formulation (Equations 2, 3 and 4) to determine the intra- and extracellular amounts (μmol) of RF+FMN and FAD in 1 L of stationary-phase cultures of *Lp* UFG9 or *Lp* B2 (Fig. 2A).

**Fig. 2.**
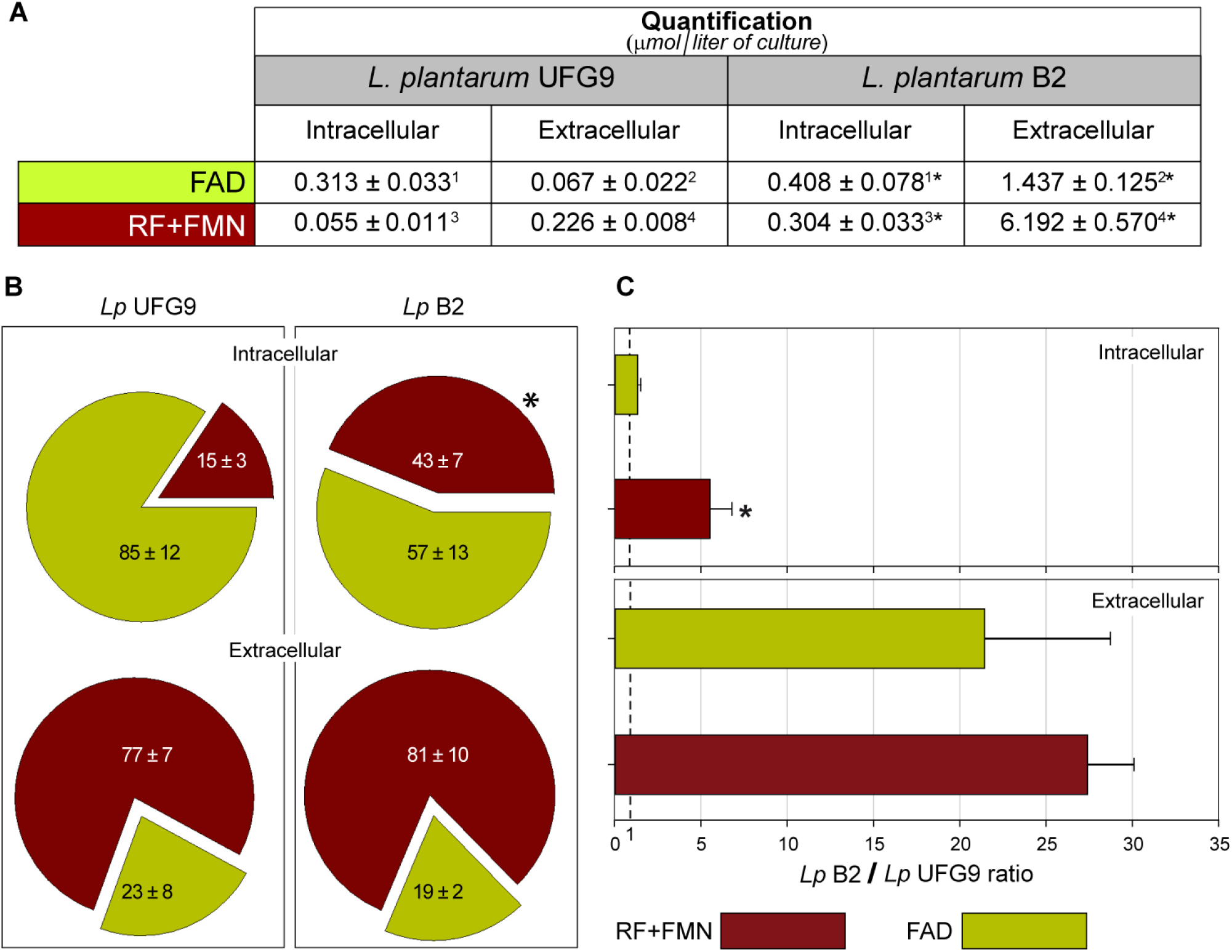
Flavins quantification. **A)** The table shows the intra- and extracellular amounts (μmol) of FAD and RF+FMN from one liter of culture at OD_480_ = 1.4 of *Lp* UFG9 and *Lp* B2. The values are expressed as mean ± SD of, at least, three independent experiments. A paired Student’s t-test was used to compare the intra- and extracellular content of each of the flavins in the *Lp* B2 strain with that of the *Lp* UFG9 strain. Asterisks denote statistically significant differences (*p* < 0.05) between pairs of values with the same superscript number. The different flavins were quantified by using the TCA-based method. **B)** Pie chart graphs showing the fraction of intra- and extracellular RF+FMN and FAD in cultures of *Lp* UFG9 and *Lp* B2. The fraction values are expressed as mean ± standard deviation (SD) of, at least, three independent experiments. The average distribution of intra- and extracellular flavins in cultures of *Lp* B2 was compared with that in *Lp* UFG9 and analyzed for statistical significant differences with a Student’s t-test. The asterisk indicates statistical significance (*p* < 0.05). **C)** Accumulation of FAD or RF+FMN in cultures of *Lp* B2 relative to *Lp* UFG9, as measured in the intra- or extracellular environment. The bar chart shows the results from, at least, three independent experiments, with the height of the bars and the horizontal lines representing the mean ratio and SD, respectively. The vertical dotted line indicates a ratio of 1. A paired t-test was used to compare the *Lp B2/Lp* UFG9 ratios of FAD and RF+FMN within the cells or in the extracellular medium, separately. The asterisk indicates statistical significance (*p* < 0.05).

FAD clearly predominated within the *Lp* UFG9 cells, where it represented ~85% of the total flavin compounds (Fig. 2B). Compared with the WT cells, the *Lp* B2 cells contained a similar concentration of FAD, but a ~6-fold higher amount of RF+FMN, which reached ~40% of the total intracellular flavin compounds in the mutant strain (Fig. 2B). On the other hand, flavin secretion was dramatically increased in *Lp* B2, and the extracellular concentrations of FAD and RF+FMN in cultures of the mutant strain were more than 20-fold higher than in the cultures of the *Lp* UFG9 WT strain (Fig. 2A and C). The relative amount of secreted flavins was notwithstanding similar for both strains, with RF and/or FMN representing ~80% of the total extracellular flavin molecules (Fig. 2B).

The intracellular concentration of flavins was estimated from the amounts of RF+FMN and FAD detected within the cells from 1 L of stationary-phase bacterial cultures, by taking into account the total cell volume (see Supporting methods). The estimated FAD concentration was 19.9±2.4 μM and 28.7±6.1 μM in *Lp* UFG9 and *Lp* B2 cells, respectively; a value of the same order of magnitude as that reported for *E. coli* (McAnulty and Wood, 2014). Concentration of RF+FMN was 3.5±0.7 μM in *Lp* UFG9 cells, and 21.4±2.4 μM within the mutant *Lp* B2 cells.

### 2.3 The leader region of the *Lp* UFG9 *rib* operon mRNA contains a potential riboswitch with a typical FMN aptamer that can be affected by the B2 mutation

Inspection of the DNA sequence upstream from the initiation codon of the first gene (*ribG*) of the *Lp* UFG9 *rib* operon revealed a potential promoter that included −35 (5’-TTGACA-3’) and extended −10 (5’-TGTTTTTAT-3’) boxes separated by a 14-nt spacer. From such a promoter, transcription of the *rib* operon may start at either of the G residues located 6 or 8 nt downstream of the −10 box (Fig. 3A). The nascent leader region of the *rib* operon mRNA (nts 1 to 145; numbering starts at the upstream G) is predicted to exhibit a dynamic folding ensemble where the most stable conformations would be populated (Fig. 3B and C). One of the two thermodynamically most favorable conformations (Fig. 3C) matches quite well the consensus sequence and structure of the FMN aptamers (Fig. 3D). Although hairpins P3, P4 and P5 are conserved in both conformations, the 3’-terminal hairpins of the corresponding multifoliate secondary structures (P6 and P6’) comprise overlapping sequences, while hairpin P2 and aptamer terminal helix P1 only exist in the aptameric conformation (Fig. 3B and C). The conserved tertiary structure of the FMN aptamers has been shown to adopt a butterfly-like scaffold through long-range interactions between different regions of the characteristic six-stem junctional secondary structure, thereby conforming a specific FMN-binding pocket (Serganov et al., 2009). Binding of the FMN effector molecule induces the stabilization of the aptamer P1 helix (Di Palma et al., 2013). Mutation B2 (in *Lp* B2) changes G139 to C, and is thus expected to destabilize the P1 helix of the aptamer-like conformation without affecting the predicted non-aptameric conformation, which would hence increase its frequency in the thermodynamic ensemble (Fig. 3B and C).

**Fig. 3.**
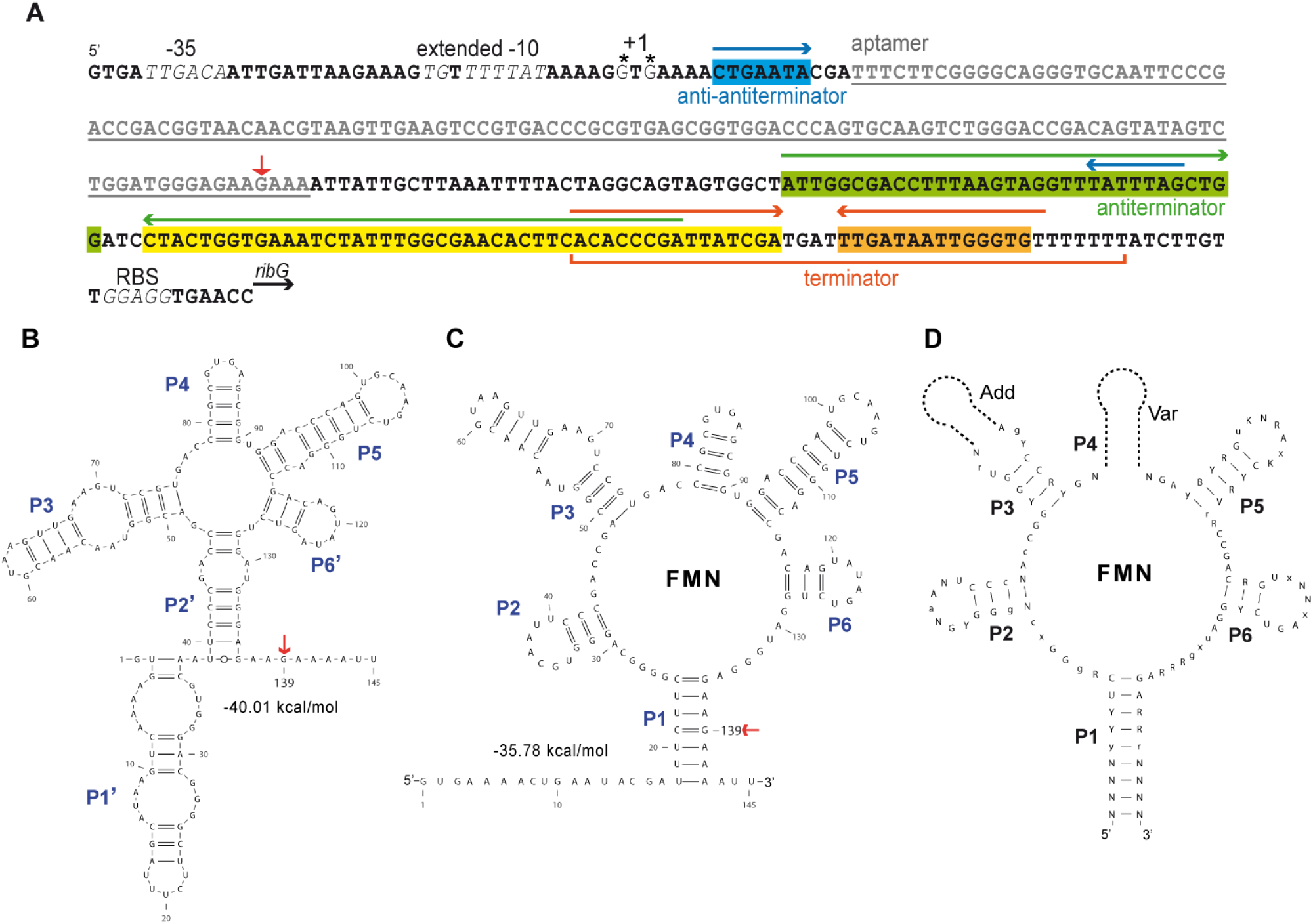
Riboswitch sequence and secondary structure prediction of the sensor domain. **A)** Sequence of the leader region of the *Lp* UFG9 *rib* operon. The −35 and extended −10 sequences of the *rib* promoter (in italic letters), two possible transcription start sites (+1, indicated by asterisks), and the ribosome binding sequence (RBS) of the first gene *(ribG)* of the operon are indicated. The figure also shows the different components of the riboswitch: aptamer (grey letters), the putative transcriptional terminator (orange bracket below the sequence, with the orange box indicating its 3’ arm), the antiterminator (green box), the anti-antiterminator (blue box), and sequences involving the 3’ arm of the antiterminator hairpin and the 5’ arm of the terminator (yellow box). The convergent arrows above the sequence indicate the complementary inverted repeats forming the anti-antiterminator (blue), antiterminator (green) and terminator (orange) secondary structures. The vertical red arrow indicates the position of the mutated base in *Lp* B2. **B** and **C)** Secondary structure predictions obtained by using *RNAfold* web server with the sequence of the nascent *rib* operon mRNA of *Lp* UFG9. The free energy associated with the two more stable structures of the aptamer domain is indicated. A red arrow in panels **B** and **C** indicates the G139 position affected by mutation B2. **D)** Conserved structure and sequence of the RFN element (scheme adapted from Vitreschak et al, 2004). Conserved regions contain invariant (uppercase) and highly conserved (lowercase) positions, although some parts of the structures are variable (Var) or facultative (Add). The conserved helices are numbered from P1 to P6, P1 being the base stem. Nucleotide notation is as follows: R, purine (A, G); Y, pyrimidine (C, T); K, keto (G, T); B, not A; V, not U; N, any nucleotide; X, not nucleotide or deletion. RNA secondary structure drawings were performed with VARNA 3.9 software (Darty et al., 2009).

A DNA region located 243 bp downstream from the proposed transcription start site could encode a putative rho-independent transcriptional terminator consisting of a hairpin structure (with a 15-bp stem and a 4-nt apical loop) followed by a 7-nt U-tract (Fig. 3A). The 283-nt region spanning from G1 to the end of the U-tract was searched for the presence of a putative riboswitch by using the *Pasific* web server (see Supporting methods). Potential key sequence and structural elements of the expression platform (anti-antiterminator, antiterminator and transcriptional terminator) were identified, and this regulatory domain was predicted to adopt two alternative configurations that corresponded to the OFF and ON states (Fig. 4A and B). In the ON state, the anti-antiterminator sequence is seized into a local 5’-terminal secondary structure (Fig. 4B), so that it cannot prevent the formation of the long antiterminator hairpin, whose 3’ arm includes bases otherwise involved in the terminator hairpin. Hence, transcription proceeds and the *rib* operon is expressed. This is expected to be the predominant state in the absence of the effector ligand, when the anti-antiterminator-seized non-aptameric conformation of the sensing domain would be highly populated (Fig. 4B). Conversely, in the OFF state, the anti-antiterminator forms a very long-range base-pairing interaction with part of the antiterminator sequence that otherwise would be involved in the antiterminator hairpin. Thus, the transcriptional terminator is generated, switching off the expression of the *rib* operon (Fig. 4A). The OFF conformation of the expression platform is predicted to predominate when the effector molecule is bound to the sensing domain of the riboswitch, since stabilization of the aptamer would leave the anti-antiterminator free to hybridize with the antiterminator sequence, leading to the generation of the terminator hairpin (Fig. 5A). In the mutant B2 riboswitch, the ON state would predominate even in the presence of the effector molecule, since the mutation would counteract the effector-induced stabilization of the aptamer terminal P1 helix.

**Fig. 4.**
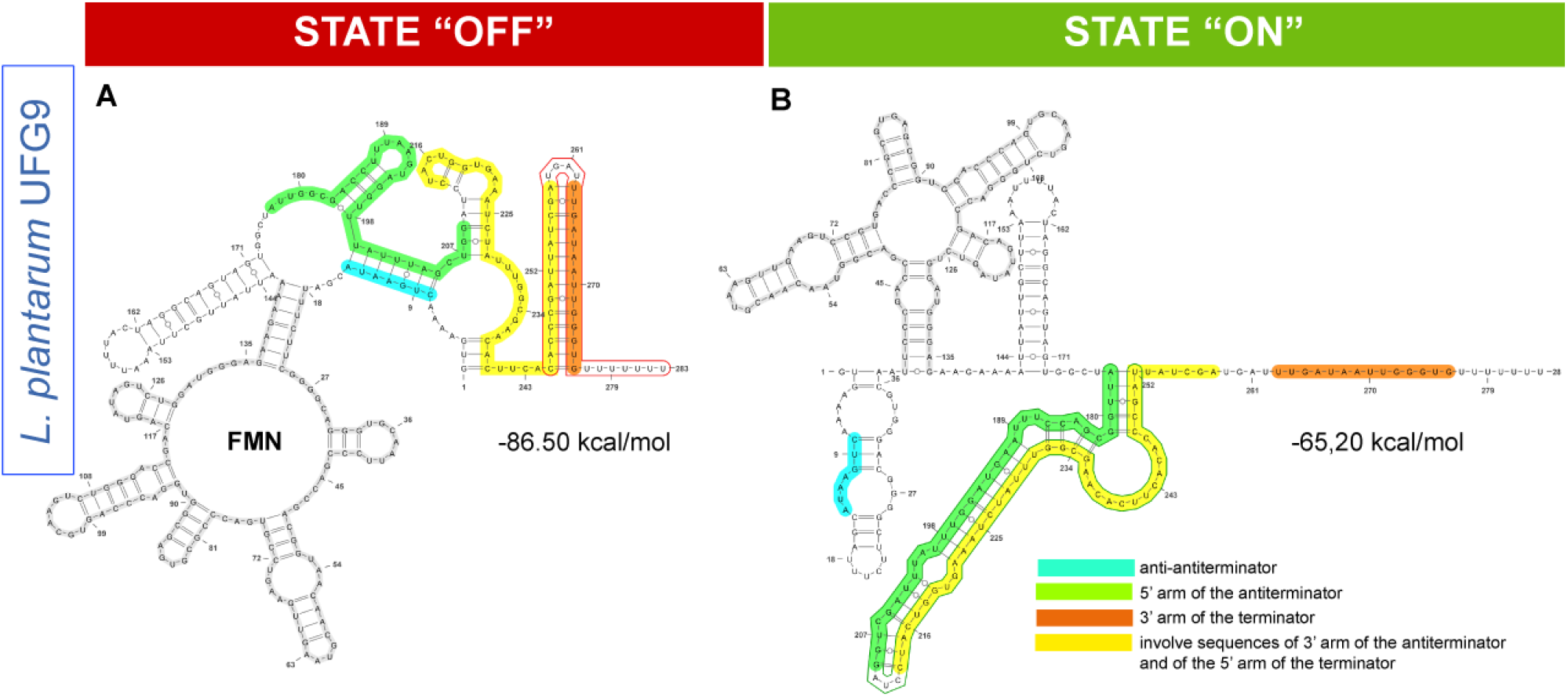
Secondary structure prediction of the regulatory domain of the riboswitch. The OFF state (**A**; terminator) was predicted by using the mFold web server. The structure of the aptamer was fixed to simulate the stabilization mediated by the ligand. The *Pasific* server predicted the ON state (**B**; antiterminator). The same color code as in Fig. 3 is used to indicate the different riboswitch elements. The free energy associated with the predicted structures is indicated in each panel.

**Fig. 5.**
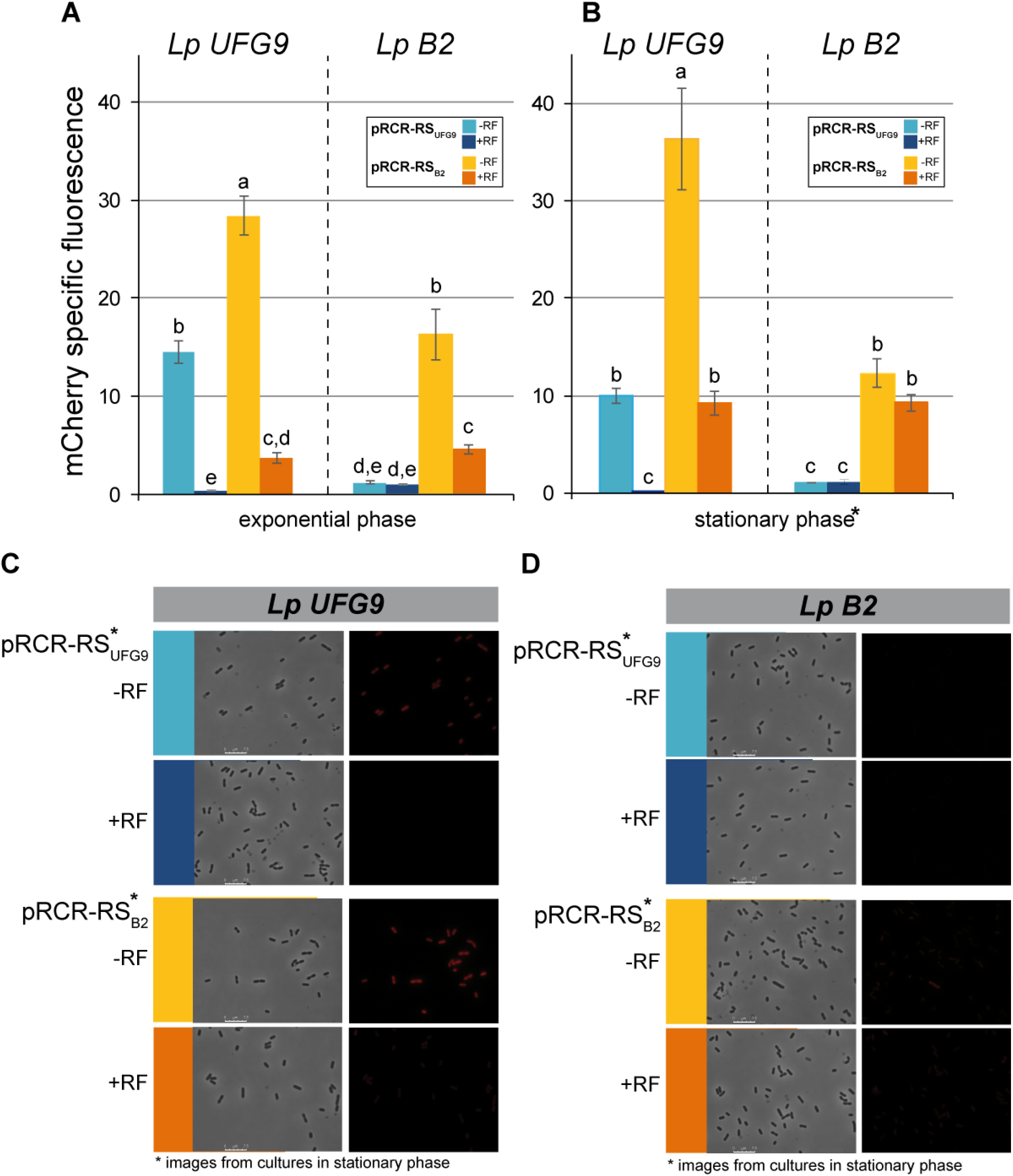
*In vivo* functional analysis of the WT and mutant B2 riboswitches. The mCherry fluorescence of Lp UFG9 and Lp B2 strains carrying pRCR-RS_UFG_ or pRCR-RS_B2_ was analyzed in cultures in exponential (**A**; OD_480_ nm = 0.7) and stationary (**B**; OD_480_ nm = 1.3) growth phase. The strains were grown in CDM medium supplemented (+RF) or not (-RF) with 1 mg/L RF. Fluorescence measurements were performed after 2 h of mCherry maturation at 30 °C in PBS. The vertical bar graphs represent the specific fluorescence (measured in the Varioskan fluorimeter) of the different strains grown in the presence or in the absence of RF. Plots display average values (bar height) and standard deviation (vertical bars) from three independent experiments. Different letters within each panel indicate significantly (p < 0.05) different mean values. Cells from cultures in stationary phase of the four strains were visualized by fluorescence microscopy (**C** and **D**). Phase contrast and fluorescence images of the same view are shown. The same color code used in **A** and **B** was used to designate the strain and the growth condition in the micrograph.

### 2.4 Mutation B2 impairs the regulatory activity of the *L. plantarum rib* operon riboswitch but does not abolishes its dependence on the effector molecule

To analyze the regulatory activity of the WT (RS_UFG9_) and B2 (RS_B2_) riboswitches, the DNA region encoding each of these elements was cloned separately in the promoter-probe vector pRCR. The recombinant plasmids (pRCR-RS_UFG9_ and pRCR-RS_B2_) were transferred to both *Lp* UFG9 and *Lp* B2, and the effector-mediated regulatory activity of the WT and B2 riboswitches was determined by monitoring the intensity of the m-Cherry fluorescence emitted when the host strains were grown to an OD_480_ of 0.7 (exponential growth phase) or 1.3 (stationary phase) in CDM either supplemented or not with 1 mg/L RF (Fig. 5). In all cases, the mutant B2 riboswitch resulted in a significantly higher expression of the mCherry-encoding *mrfp* gene than the WT riboswitch (Fig. 5), a finding that is consistent with the RF-overproducing phenotype of the *Lp* B2 strain.

Compared with the values observed in CDM lacking RF, expression regulated by the WT riboswitch decreased to ~2% for the *Lp* UFG9 host cells grown to either the exponential or the stationary phase in medium supplemented with the vitamin. A lower inhibition due to the presence of RF in the culture medium was found for the B2-riboswitch-regulated expression in the UFG9 genetic background, since the specific fluorescence values of cultures at the exponential and stationary phase were, respectively, 13% and 25% of those observed when the bacterial cells were grown in RF-lacking CDM (Fig. 5A and B). Overall, these results indicate that *Lp* UFG9 imports RF when grown in CDM supplemented with the vitamin. Within the cells, RF would be rapidly converted into FMN, which is the main effector of the FMN riboswitches as it binds to their sensing domain with higher affinity than FAD or RF (Winkler et al., 2002). The results also show that the mutant B2 riboswitch, although severely impaired in its regulatory activity, does not drive constitutive expression of the target gene/operon but still remains responsive to the presence of the effector.

The levels of WT- or B2-riboswitch-regulated *mrfp* expression in the *Lp* B2 host grown to either exponential or stationary phase in RF-lacking CDM were significantly (*p* < 0.05) decreased compared with those observed in the UFG9 (WT) genetic background (Fig. 5A and B), which is consistent with the higher intracellular concentration of RF+FMN in the mutant cells (Fig. 2A and C and Table 1). When the *Lp* B2 host cells were grown to exponential or stationary phase in RF-supplemented medium, the level of *mrfp* gene expression regulated by the B2 riboswitch decreased, respectively, to ~30% (statistically significant) and ~75% (not statistically significant) of the values observed when the bacterial host was grown in RF-free CDM (Fig. 5A and B). Compared with the mCherry fluorescence levels observed in CDM, no further decrease of the *mrfp* expression regulated by the WT riboswitch occurred when the *Lp* B2 host cells were grown in CDM plus RF, and near basal values were obtained irrespective of whether the vitamin was added or not to the culture medium (Fig. 5A and B). This result suggests that the high intracellular concentration of RF+FMN in *Lp* B2 provides almost complete inhibition of the WT-riboswitch-regulated gene expression.

**Table 1.**
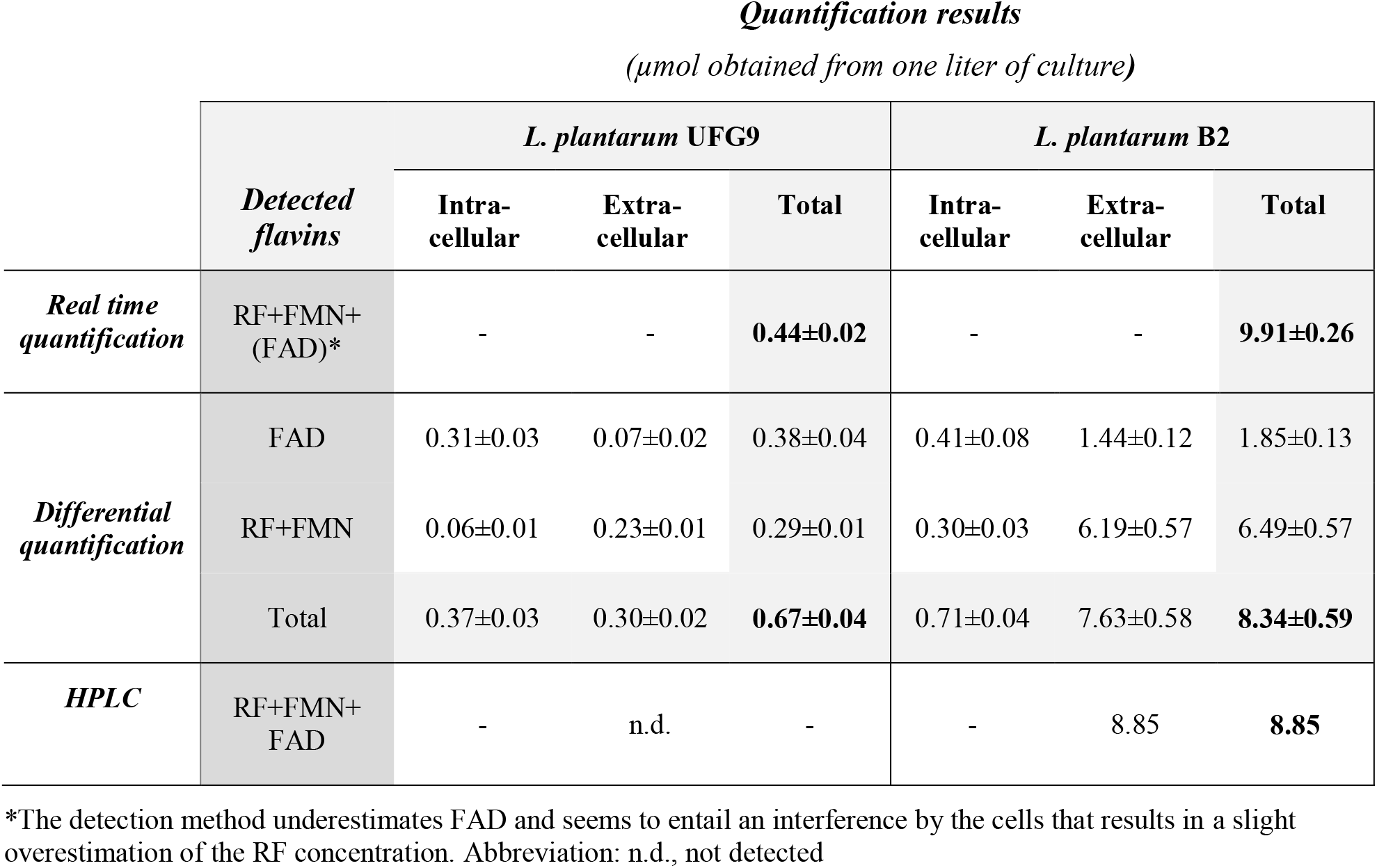
Summary of flavins quantification by different methods.

Fluorescence microscopy was also used to qualitatively analyze the WT- or B2-riboswitch-regulated expression of the *mrfp* gene in individual cells with either UFG9 or B2 genetic background grown to the stationary phase (Fig. 5C and D). A quite homogeneous distribution of the fluorescence was observed among the cells of each of the groups studied, and the results agreed with those obtained in the quantitative analysis of the intensity of the mCherry fluorescence in the corresponding bacterial cultures (Fig. 5A and B). The highest level of fluorescence was observed in cells of *Lp* UFG9/pRCR-RS_B2_ (carrying the *mrfp* gene under regulation by RS_B2_) grown in RF-free CDM. A lower level of fluorescence was observed in cells of *Lp* UFG9/pRCR-RS_UFG9_ (harboring the *mrfp* gene under regulation by RS_UFG9_) grown in RF-free CDM, *Lp* UFG9/pRCR-RS_B2_ grown in CDM supplemented with RF, and *Lp* B2/pRCR-RS_B2_ grown in medium either supplemented or not with the vitamin. No fluorescence could be detected in cells of *Lp* UFG9/pRCR-RS_UFG9_ grown in CDM supplemented with RF and of *Lp* B2/pRCR-RS_UFG9_ grown in either RF-free or RF-containing CDM (Fig. 5C and D).

## 3 Discussion

*Lp* UFG9 is a natural RF-producing LAB strain that was isolated from traditional Italian wheat sourdough, while the RF-overproducing *Lp* B2 strain was obtained by selection for resistance against roseoflavin (Arena et al., 2014). Here, these two isogenic strains have been compared with respect to their capacity for flavin production, their intracellular and secreted flavin profiles, and the regulatory activity of their *rib* operon riboswitch. Two main objectives were pursued: i) to identify and quantify, for the first time, the intra- and extracellular flavin compounds produced by a natural isolate of *L. plantarum* and one roseoflavin-resistant derivative in order to gain information on the bioavailability of the vitamin B_2_ synthesized by these LAB, and ii) to analyze the effect of the B2 mutation on the regulation of the *rib* operon expression in order to understand the molecular bases underlying one of the highest riboflavin yields by a LAB strain reported so far.

Our studies have revealed that FAD is clearly the predominant flavin within the WT *Lp* UFG9 cells, and that its concentration remains basically unchanged in the *Lp* B2 mutant cells (Fig. 2A and C). The estimated intracellular concentrations of RF+FMN and FAD in *Lp* UFG9 are ~3.5 μM and ~20 μM, respectively, which do not greatly differ from the values reported for the Firmicute bacterium *Amphibacillus xylanus* (Kimata et al., 2018). In contrast, the intracellular concentration of RF+FMN in *Lp* B2 was estimated to be ~20 μM. As far as we know, this is a unique information on the intracellular concentration of flavins in LAB. On the other hand, the biosynthesized RF (and/or FMN) was mostly secreted by both *Lp* UFG9 and *Lp* B2 cells (Fig. 2B), and its extracellular concentration almost reached 5 μM in stationary-phase *Lp* B2 cultures (Fig. 2A). To the best of our knowledge, no other analyses on the distinct flavin secretion profile in LAB have been reported so far. For a number of LAB strains, however, the total concentration of flavins in the medium or in the entire culture has been determined by HPLC after their acid-mediated conversion into RF (Burgess et al., 2006; Juarez del Valle et al., 2014; Russo et al., 2016; Ge et al., 2020). Here, we have employed three different methods to quantify the flavins produced by the UFG9 and B2 strains of *L. plantarum* (Table 1). Considering the location of the flavins detected and the sensitivity to the different flavin compounds, the results obtained by all three methods were found to be quite consistent (Table 1). Compared with other RF-overproducing LAB strains, *Lp* B2 renders the highest amount of extracellular RF+FMN, as quantified by either real-time fluorometric determination in the Varioskan or fluorometry subsequent to HPLC (Burgess et al., 2004, 2006; Capozzi et al., 2011; Arena et al., 2014; Juarez del Valle et al., 2014; Russo et al., 2014a; Ge et al., 2020).

The use of RF-overproducing LAB, like *Lp* B2, is most suitable for the manufacturing of vitamin B2-enriched processed foods as it allows omitting additional fortification steps, hence decreasing the production costs (Thakur et al., 2016). Moreover, these bacteria can improve the quality and safety of the food product by i) contributing to the fermentation process, ii) conferring functional and nutritional properties other than RF supply, and iii) exerting protective effect against foodborne pathogens (Capozzi et al., 2011; Russo et al., 2014a, 2014b, 2015b; Zhu et al., 2020; Yepez et al., 2019). *Lp* B2 seems to display a great metabolic versatility that allows its use for the manufacturing of a variety of vitamin B_2_-fortified foods prepared from fruit and vegetable matrices (Capozzi et al., 2011; Russo et al., 2014a, 2015b, 2016; Zhu et al., 2020). As shown in Fig. 2A, the flavins secreted by *Lp* B2 mainly consist of RF (and/or FMN) (~80%), which is then the compound in charge of the vitamin B2 biofortification of foods by this bacterium. In contrast, the major dietary source of vitamin B_2_ is FAD, as this is the prevalent flavin compound within both eukaryotic and bacterial cells (Kimata et al., 2018; Kang et al., 2019; Fig. 2). Absorption across the intestinal mucosa requires the enzymatic hydrolysis of FAD to RF, which is inhibited in the presence ethanol and its major metabolite, acetaldehyde, a condition matched in alcohol abusers among whom a high prevalence of ariboflavinosis is well documented (Pinto et al., 1987). Thus, RF displays a higher bioavailability than FAD, even in the presence of ethanol and acetaldehyde, which entails an additional benefit of using *Lp* B2 (as well as other RF-overproducing LAB strains) for the manufacturing of vitamin B_2_-fortified foods and/or probiotics that may have positive clinical implications for nutritional treatment of specific population groups. As a second main objective of the present work, we have analyzed the regulatory activity of the potential riboswitch spanning the 5’-untranslated region of the *Lp* UFG9 *rib* operon mRNA, as well as the effect of mutation B2 on the riboswitch functionality (Figs. 3, 4 and 5).

A model for the regulatory mechanism of the *Lp* UFG9 *rib* operon riboswitch has been proposed that combines the ON or OFF state of its regulatory domain (as predicted by the *Pasific* web server) with the most stable (according to the *RNAfold* web server) non-aptameric or aptameric conformation of its sensing domain, respectively (Fig. 4). The model predicts that the riboswitch ON state, characterized by a long antiterminator hairpin, will be populated in the absence of the effector molecule due to the seizing of the anti-antiterminator sequence in a 5’-terminal secondary structure of the non-aptameric conformation of the sensing domain. Conversely, binding of the effector to the aptamer will stabilize the long-range P1 helix, leaving the anti-antiterminator sequence free to hybridize with the antiterminator, thus precluding the antiterminator hairpin and leading to the generation of the transcriptional terminator structure, which characterizes the OFF state of the riboswitch (Fig. 4). Mutation B2 would destabilize the aptamer terminal P1 helix, hence counteracting the effect of ligand binding (Di Palma et al., 2013). Analysis of the *in vivo* regulatory functionality of the *L. plantarum rib* operon WT and mutant B2 riboswitches shows that: i) mutation B2 results in an increased expression of the *rib* operon, irrespective of whether the culture medium was supplemented or not with RF; ii) the greatly impaired regulatory activity of the mutant riboswitch is the most likely cause of the flavin overproduction phenotype of *Lp* B2, since no other mutation was found in the leader region of the *rib* operon; and iii) despite its impaired functionality, the mutant B2 riboswitch still remains responsive to the effector molecule (Fig. 5). This latter observation indicates that, although with a lower frequency than in the WT riboswitch, the aptameric conformation of the B2 riboswitch sensing domain can still occur, and its P1 terminal helix can become stabilized by binding of the effector, thus leading to the OFF state.

The model presented here for the riboswitch of the *Lp* UFG9 *rib* operon includes, among other key structural elements, an antiterminator hairpin formed by short-range base pairing (Fig. 4). This contrasts with the overall schemes proposed for the *rib* operon riboswitch of *B. subtilis* (Abbas and Sibirny, 2011) and *L. lactis* (Burgess et al., 2004), where an antiterminator helix is generated by long-range base pairing (Fig. 6). Verification of any of the proposed models will require *in vivo* and/or *in vitro* probing of the entire riboswitch RNA structure both in the absence and in the presence of the effector. It is worth mentioning that one of the *L. lactis* roseoflavin-resistant mutants that produces highest amounts of RF has been reported to contain a G→U point mutation in the anti-antiterminator sequence located at the 3’-end of the RFN element ((Burgess et al., 2004); Fig. 6). In the *L. lactis* riboswitch model, the anti-antiterminator overlaps with the sequence of the 3’-arm of the aptamer P1 helix, and a visual inspection of this stem reveals that the residue affected by the mutation is equivalent to the G139 residue that is changed to C in the mutant riboswitch of *Lp* B2 (Fig. 6). Despite this equivalence, the regulatory behavior seems to differ between the *L. lactis* and *L. plantarum* mutant riboswitches. In both bacteria, the *in vivo* regulatory activity of both the WT and the mutant riboswitch was analyzed by monitoring the expression of a plasmid-borne reporter gene encoding either β-Galactosidase (in *L. lactis;* (Burgess et al., 2004)) or mCherry (in *L. plantarum;* Fig. 5). A more complete picture can be drawn for the *L. plantarum* regulatory element since the WT and B2 riboswitches were analyzed in both genetic backgrounds in the present work, whereas the *L. lactis* WT and mutant riboswitches were only tested in their cognate background. The *L. lactis* mutant riboswitch drove similar (constitutive) expression of the reporter gene in the mutant genetic background, irrespective of the presence or absence of RF in the culture medium, although the β-Galactosidase activity was quite lower than that obtained, in the absence of RF, from the WT riboswitch in the WT genetic background (Burgess et al., 2004). In contrast, the *Lp* B2 riboswitch showed an effector-responsive behavior in both the WT and the B2 genetic backgrounds, and the levels of mCherry fluorescence were similar to or higher than those observed for the WT riboswitch in either background, when the bacterial hosts were grown in RF-free medium. The distinct behavior of the *L. lactis* and *L. plantarum* riboswitches carrying a point mutation in an equivalent G residue of the potential P1 helix can be due to the high intracellular concentration of the FMN effector causing saturation of the aptamer in the lactococcal cells (irrespective of whether they were grown in the absence or presence of RF). In contrast, although the high intracellular concentration of the effector in the *Lp* B2 cells causes a decreased expression of the reporter gene relative to the WT cells, the mutant B2 riboswitch is still sensitive to an increase of the effector concentration resulting from the presence of RF in the culture medium. The effector-responsive regulatory activity of the B2 riboswitch is most clearly observed in the WT cells, a condition that was not tested for the *L. lactis* mutant riboswitch.

**Fig. 6.**
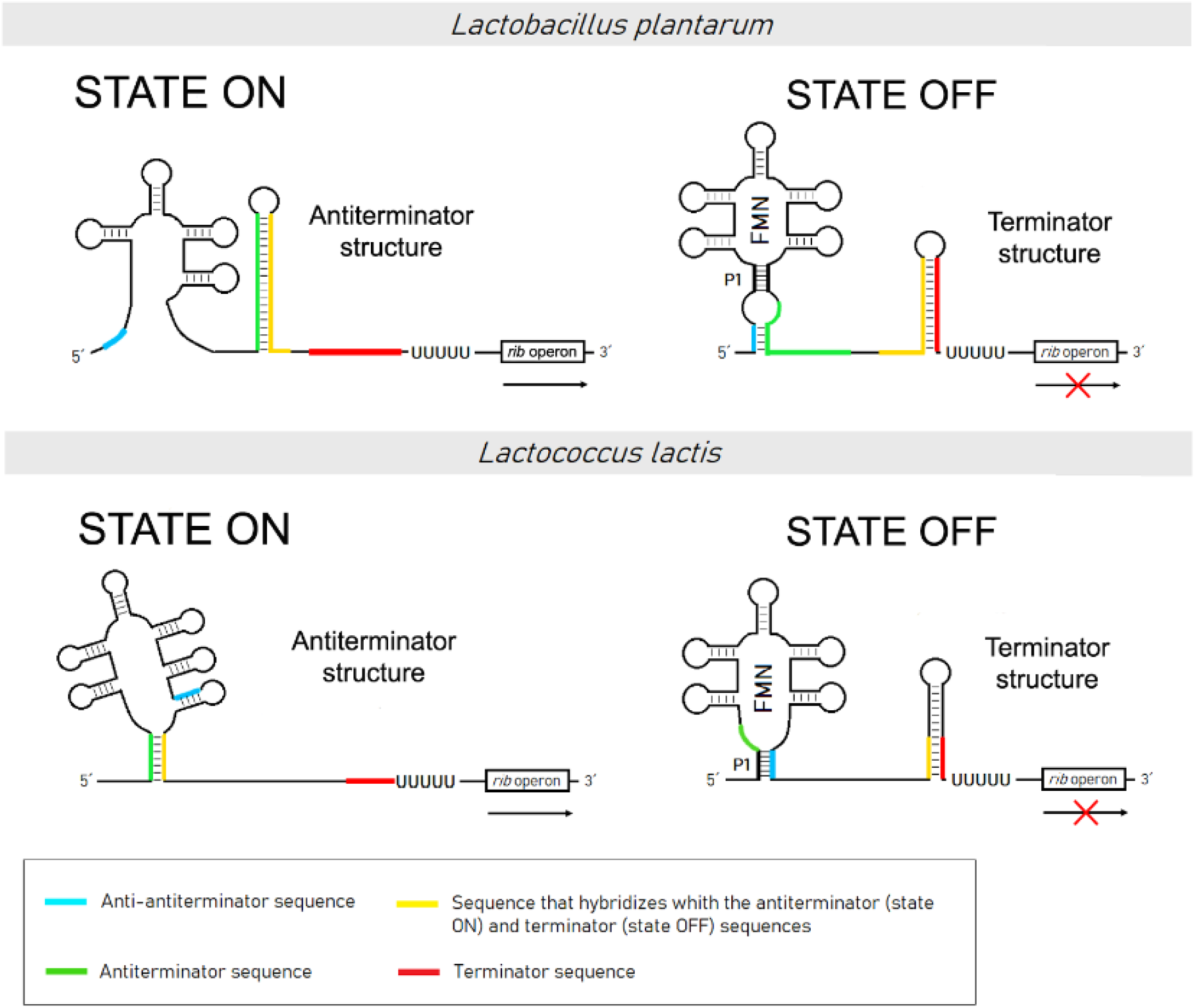
Mechanism proposed for the riboswitch-mediated *rib* operon regulation in *Lp* UFG9 and *L. lactis*. The structural elements indicated in the figure include the antiterminator hairpin, the transcriptional terminator and the aptamer. The putative antiterminator structure (state ON) is disrupted after binding of FMN and formation of the aptamer P1 helix (state OFF).

Although the results obtained in the present work are compatible with the proposed model of the *L. plantarum rib* operon riboswitch, validation of this model will require not only probing of the secondary structure of the mRNA leader region, but also analyzing the effect of deletions of specific elements (such as the anti-antiterminator, antiterminator and terminator) on the regulatory capacity of the riboswitch. Curiously enough, the analysis of roseoflavin-resistant spontaneous mutants isolated from LAB shows that deletions in the putative RFN element result in a phenotype of similar or even lower overproduction of RF compared with some point mutations in this element, while deletions or insertions affecting the transcriptional terminator of the expression platform of the riboswitch give rise to phenotypes of highest RF overproduction (Burgess et al., 2004, 2006).

The extraordinary capacity of *Lp* B2 to produce and secrete RF, characterized in the present work, together with its metabolic versatility (Capozzi et al., 2011; Russo et al., 2014a, 2014b, 2015b; Zhu et al., 2020) and its probiotic potential (Arena et al., 2014; Russo et al., 2015a) make this strain very useful for the manufacturing and preserving of vitamin B_2_-enriched fermented and otherwise processed foods, which additionally may serve as delivery vehicles for the bacterium. Since changes in the gut microbiota composition can severely affect our dietary B-vitamin requirements (Magnúsdóttir et al., 2015), gut colonization by the RF-overproducer *Lp* B2 strain could allow the human host and their intestinal microbiota to benefit from both its ability to produce and secret the vitamin and its probiotic properties.

## 4 Experimental procedures

### 4.1 Bacterial strains and growth conditions

*Lp* UFG9 and *Lp* B2, the latter deposited to the Spanish Type Culture Collection (CECT, Paterna, Spain) under the code CECT 8328 (Arena et al., 2014), were routinely cultivated in MRS broth at 30 °C. To evaluate the RF production and to analyze the functionality of the RS, these strains were cultivated in chemically defined medium (CDM; Table S2) as reported by Russo et al., 2014a. The *L. plantarum* strains carrying the constructions based on the pRCR plasmid were cultivated in MRS broth containing 20 μg/mL chloramphenicol (Cm). *Streptococcus pneumoniae* 708 (Lacks and Greenberg, 1977) was cultivated at 37 °C without aeration in AGCH medium (Lacks, 1968) supplemented with 0.3% sucrose and 0.2% yeast extract. *Escherichia coli* DH5α harboring vector pRCR was cultivated at 37 °C with aeration in lysogeny broth (LB) medium. Solid media were prepared from the corresponding liquid broth by adding agar to a final concentration of 1.5%.

### 4.2 Plasmid and genomic DNA preparation

Plasmid pRCR, a promoter-probe vector carrying a promoterless *mrfp* gene (Mohedano et al., 2014), was isolated from *E. coli* DH5α and purified by using the Favorprep Plasmid Extraction Mini kit (Favorgen). Plasmids pRCR-RS_UFG9_ and pRCR-RS_B2_ were purified from *S. pneumoniae* 708 as described (Ruiz-Cruz et al., 2010). Genomic DNA (gDNA) from *L. plantarum*, used as template for PCR, was isolated by using the Wizard Genomic DNA Purification kit (Promega). Prior to the lysis step, collected cells were resuspended in a solution containing 50 mM EDTA, 30 mg/mL lysozyme and 25 U of mutanolysin, and incubated at 37 °C for 40 min.

DNA integrity was checked by 0.8%-agarose gel electrophoresis followed by staining with GelRed (Biotium). Concentration of the DNA was determined with a Qubit fluorometer by using the Qubit HS dsDNA Assay kit (Molecular Probes).

### 4.3 Plasmid constructions

To construct plasmids pRCR-RS_UFG9_ and pRCR-RS_B2_, DNA from pRCR was digested with XbaI, which generates 5’-P protruding ends, and with SmaI, which leaves 5’-P blunt ends. DNA encoding the entire WT or mutant *rib* operon riboswitch (including the potential promoter and transcriptional terminator sequences) was PCR-amplified by using specific primers ForRFN and RevRFN (Table S1), and gDNA from *Lp* UFG9 or *Lp* B2, respectively, as template. EcoRV and XbaI restriction sites were incorporated at the 5’ end of the ForRFN and RevRFN primers, respectively. The resultant RS_UFG9_ and RS_B2_ fragments (392 bp) were purified with the QIAquick PCR purification kit (Qiagen), and subsequently subjected to sequential digestion with XbaI and EcoRV (which also leaves 5’-P blunt ends). After digestion, both the PCR fragments and the vector were purified with the QIAquick PCR purification kit (Qiagen) and ligated with the T4 DNA ligase (New England Biolabs) at an insert to vector molar ratio of 3:1. To reduce the presence of recircularized pRCR vector, the ligation mixture was digested with SmaI, whose recognition sequence is missing in the recombinant constructions. Finally, the digested ligation mixture was used to transform *S. pneumoniae* 708 competent cells. This pneumococcal strain was used to obtain the recombinant plasmid constructs due to its high efficiency of natural transformation. The correct nucleotide sequence of the two constructs was confirmed by automated DNA sequencing. The recombinant plasmid constructs were isolated from *S. pneumoniae* and then introduced by electrotransformation in *L. plantarum*.

### 4.4 Transformation of bacterial species with plasmid DNA

Competent cells of *S. pneumoniae* 708 were prepared and transformed as described (López et al., 1982). Transformants were selected with 3 μg/mL Cm. For transfer of pRCR-RS_UFG9_ and pRCR-RS_B2_ to *L. plantarum* strains, 2.5 μg of the plasmid DNAs were used for electroporation (25 μF, 1.3 kV, and 200 Ω, in 0.1-cm cuvettes) as previously described (Berthier et al., 1996). Transformant cells were recovered in 0.5 mL of MRS supplemented with additional 1% glucose and 80 mM MgCl_2_, and incubated 2 h at 30 °C without aeration. Expression of the plasmidic Cm^R^ determinant (the *cat* gene) was induced with Cm 0.5 μg/mL within the last 30 min of incubation. Transformants were selected in MRS-agar plates containing 20 μg/mL Cm. Cells carrying the desired constructions were identified by colony PCR. To this end, a small portion of a transformant colony was resuspended in 100 μL of PBS 1X, and 0.5 μL of this suspension was used as template in the PCR. The reaction was performed in the presence of 1.5 mM MgCl_2_, 25 pmol of each primer (STOPCatFor y RevCherry; Table S1), 0.2 mM dNTPs and 0.5 U of Taq DNA polymerase. Thermal cycling conditions were as follows: initial denaturation at 94 °C for 3 min, followed by 30 cycles of 94 °C for 30 s (denaturation), 59 °C for 30 s (primer annealing), and 72 °C for 3 min (elongation). The size of the amplified fragments was analyzed by 1%-agarose gel electrophoresis and DNA staining with GelRed (Biotium), and the correct insert’s nucleotide sequence was confirmed by automated DNA sequencing. Additionally, the sequence of the chromosomal *rib* operon riboswitch of the *L. plantarum* strains harboring pRCR-RS_UFG9_ or pRCR-RS_B2_ was checked. For this purpose, the chromosomal riboswitch sequences were PCR-amplified by using specific primers FUPRSLp and RevRFN (Table S1), and gDNA from the mentioned strains as template. Primer FUPRSLp allows specific amplification of the chromosomal sequences, as it hybridizes with a region located upstream of the cloned sequence. The resultant PCR fragments were purified and subjected to automated sequencing.

### 4.5 Flavin quantification

#### 4.5.1 Real time quantification of riboflavin production

Simultaneous measurement of cell growth and RF fluorescence was performed essentially as described (Mohedano et al., 2019). *L. plantarum* cells were grown in MRS at 30 °C to an OD_600_ of 0.5, washed twice with PBS, and diluted 10 times in CDM. Then, 200-μL triplicates of the cultures were loaded in a 96-well optical bottom plate, incubated at 30 °C and analyzed during growth in real time in a Varioskan equipment. Cell density (absorbance at 480 nm), and RF fluorescence (λ_ex/em_ 450/530 nm; Massey, 2000) were measured at 30 min intervals. To ascertain the intra- or extracellular location of the real-time detected flavins, two 600-μL aliquots of stationary-phase *L. plantarum* cultures were centrifuged, and the supernatants collected. The cellular pellets were washed twice with PBS 1X and resuspended in the same volume of either fresh CDM or their own collected supernatant. Finally, 200-μL triplicates of both cell suspensions, as well as of the supernatants and uncentrifuged cultures, were loaded in a 96-well optical bottom plate and analyzed for RF fluorescence as above.

The bacterial growth rate (μ) was calculated during the exponential growth phase as indicated in (Garay-Novillo et al., 2019). To quantify RF production, a calibration curve was performed with solutions of RF in CDM at 0.25, 0.5, 1.25, 2.5 and 5 mg/L. Triplicate 200-μL aliquots of each solution were transferred to a 96-well optical bottom plate and analyzed for RF fluorescence in a Varioskan fluorimeter. Prior to data analysis, fluorescence of a CDM blank solution was subtracted. Then, the corrected fluorescence values were plotted against the concentrations of RF (mg/L), and the experimental data were fitted to a lineal regression model of equation:

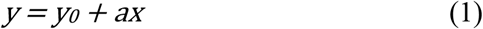

where *y* represents fluorescence of RF (arbitrary units), *y_0_* is the intercept of the regression line with the y-axis, *x* is the concentration (mg/L) of RF, and *a* is the slope of the linear regression fit (i.e., the increase in fluorescence for each 1 mg/L increase in the RF concentration), which was assigned as the intrinsic RF fluorescence.

#### 4.5.2 Differential quantification of flavins

A method for the differential quantification of RF+FMN and FAD accumulated inside the cells or secreted to the extracellular medium was developed. This method was based on the conversion of FAD into FMN by acid treatment at 37 °C and on the different intrinsic molar fluorescence of FAD and FMN or RF (Chen et al., 2013).

*L. plantarum* cells were pre-grown in MRS and subcultured in CDM as depicted in the previous section. The cellular pellet and the supernatant of early-stationary-phase cultures were analyzed separately. For determination of intracellular flavins, the cell pellet from a 250-μL aliquot was washed twice with 0.9% NaCl and resuspended in 1.5 mL of 10% trichloroacetic acid (TCA) and incubated on ice for 60 min. Then, the sample was centrifuged at 34500 × *g* at 4 °C for 6 min and the supernatant transferred to a clean tube. To analyze the extracellular flavin content, 300 μL of 50% TCA was added to the supernatant from 1.2 mL of culture, and the mixture was incubated on ice for 10 min. A first 600-μL aliquot of the samples from the pellet and the supernatant was immediately neutralized by adding 150 μL of 4 M K_2_HPO_4_, while a second 600-μL aliquot was incubated away from light at 37 °C for 16 h to get the complete hydrolysis of FAD to FMN, and then neutralized by adding 150 μL of 4 M K_2_HPO_4_. The fluorescence (λ_ex/em_ 450/530 nm) of the different samples was measured in the Varioskan fluorimeter as indicated above.

Calibration curves were elaborated with RF and FAD solutions of defined concentrations prepared in media simulating the neutralized samples from the pellet and the supernatant of the cultures (Fig. S1). Triplicate 200-μL aliquots of each solution were transferred to a 96-well optical bottom plate and analyzed for fluorescence (λ_ex/em_ 450/530 nm). The corrected fluorescence was related to the total concentrations of RF or FAD following a lineal equation. The intrinsic molar fluorescence of RF or FMN (f_RF_) and of FAD (f_FAD_) calculated from the calibration curve for the extracellular samples was 8.38±0.30 and 1.33±0.03 μM^-1^, respectively. In the case of the calibration curve for the intracellular samples, the values of f_RF_ and f_FAD_ were 9.65±0.55 and 1.88±0.11 μM^-1^, respectively. The flavin content was calculated by using the following equations:

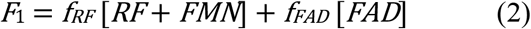

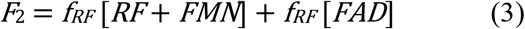

where *F_1_* and *F_2_* are the fluorescences of the samples neutralized immediately and after 16 h of incubation at 37°C, respectively. The FAD concentration was calculated from equation:

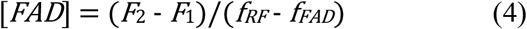

### 4.6 Analysis of the expression of the *mrfp* gene in *L. plantarum*

To analyze the mCherry fluorescence emitted by the *L. plantarum* strains harboring pRCR-RS_UFG9_ or pRCR-RS_B2_, cells grown in MRS with 20 μg/mL Cm were transferred to Cm-containing CDM supplemented or not with 1 mg/L of RF, as described above. Aliquots of the different cultures were taken at OD_480_ of 0.7 and 1.3, and the cells were harvested, washed with PBS 1X, concentrated 5-fold in the same buffer, and incubated for 2 h at 30 °C to allow maturation of the mCherry chromophore (Garay-Novillo et al., 2019). Finally, triplicate 200-μL aliquots of each bacterial cell suspension were transferred to a 96-well optical bottom plate and analyzed for mCherry fluorescence (λ_ex/em_ 587/612 nm) in a Varioskan fluorimeter.

### 4.7 Statistical analysis

Statistical analyses were carried out using SigmaPlot software (v12.5, Systat software, San Jose, CA, United States). Differences in the intra- and extracellular flavin content between *Lp* UFG9 and *Lp* B2 were analyzed by Student’s t-tests. Differences in mCherry fluorescence emitted by the *L. plantarum* strains carrying pRCR-RSUFG9 or pRCR-RSB2 were analyzed by one-way ANOVA. The level of significance in both cases was set at *p* < 0.05.

## 5 Acknowledgments

This work is based upon the work from COST Action 18101 SOURDOMICS – Sourdough biotechnology network towards novel, healthier and sustainable food and bioprocesses (https://sourdomics.com/; https://www.cost.eu/actions/CA18101/), where GS and JAR-M are members of the working group 4. Pasquale Russo is the beneficiary of a grant by MIUR in the framework of ‘AIM: Attraction and International Mobility’ (PON R&I2014-2020) (practice code D74I18000190001). We thank Dr. Paloma López and Dr. Mari Luz Mohedano for providing valuable information. We thank Ivan Belinchón-Esteban, Erika Cabanillas-Oscco and Francisca Álvarez-Juárez for critical reading of the manuscript.

## 8 Author contributions

GS conceived and designed the research. IR and JAR-M carried out the experiments and prepared the figures. NDS contributed to the flavin quantification experiments. JAR-M contributed to the writing of Experimental Procedures section. GS wrote the manuscript with support from PR and GS. PR and GS provided the *L. plantarum* UFG9 and B2 strains. All authors discussed the results and contributed to the final manuscript.

## Supporting information

### 12 Supporting Methods

#### 12.1 *In silico* analysis of the riboswitches

The aptamer domain was predicted by RNAfold web server (Lorenz et al., 2011) which has several options of folding conditions and enables the study of the conformation of the nascent RNA. In these analysis, the Turner 2004 model was applied (Mathews et al., 2004). The conformation of the complete riboswitch in its “ON” state was predicted by the *Pasific* server (Millman et al., 2017). The regulatory element of the riboswitch in the “OFF” state was predicted by mFold web server (Zuker, 2003).

#### 12.2 Fluorescence microscopy

Cells of stationary-phase cultures of *L. plantarum* strains harboring pRCR-RS_UFG9_ or pRCR-RS_B2_ were sedimented by centrifugation and resuspended in PBS 1X to obtain a ten-fold concentrated bacterial suspension, from which a10-μL aliquot was used, without fixing, for phase contrast and fluorescence microscopy analysis as depicted in (Mohedano et al., 2019).

#### 12.3 Cell volume determination

*L. plantarum* cells were pre-grown in MRS and subcultured in CDM to the early-stationary-phase. The cultures were then diluted 1:500 or 1:1000 directly in PBS 1X, and 100 μl of the dilutions were mixed by inversion with 10 ml of CASY^®^ton in a CASY^®^cup to get the final bacterial suspension for the measurement of the cell volume. The prepared cell suspension was placed under the measuring 60 μm capillary. The CASY^®^device measured three aliquots of 200 μl and the software calculated the average cell volume per milliliter. Based on this parameter and taking into account the cell culture dilutions performed, we calculated an average cell volume per liter of stationary-phase *L. plantarum* culture of 15.73 ± 0.45 ml.

### 13 Supporting Figures and Tables

#### 13.1 Supporting Figures

**Supporting Fig. S1.**
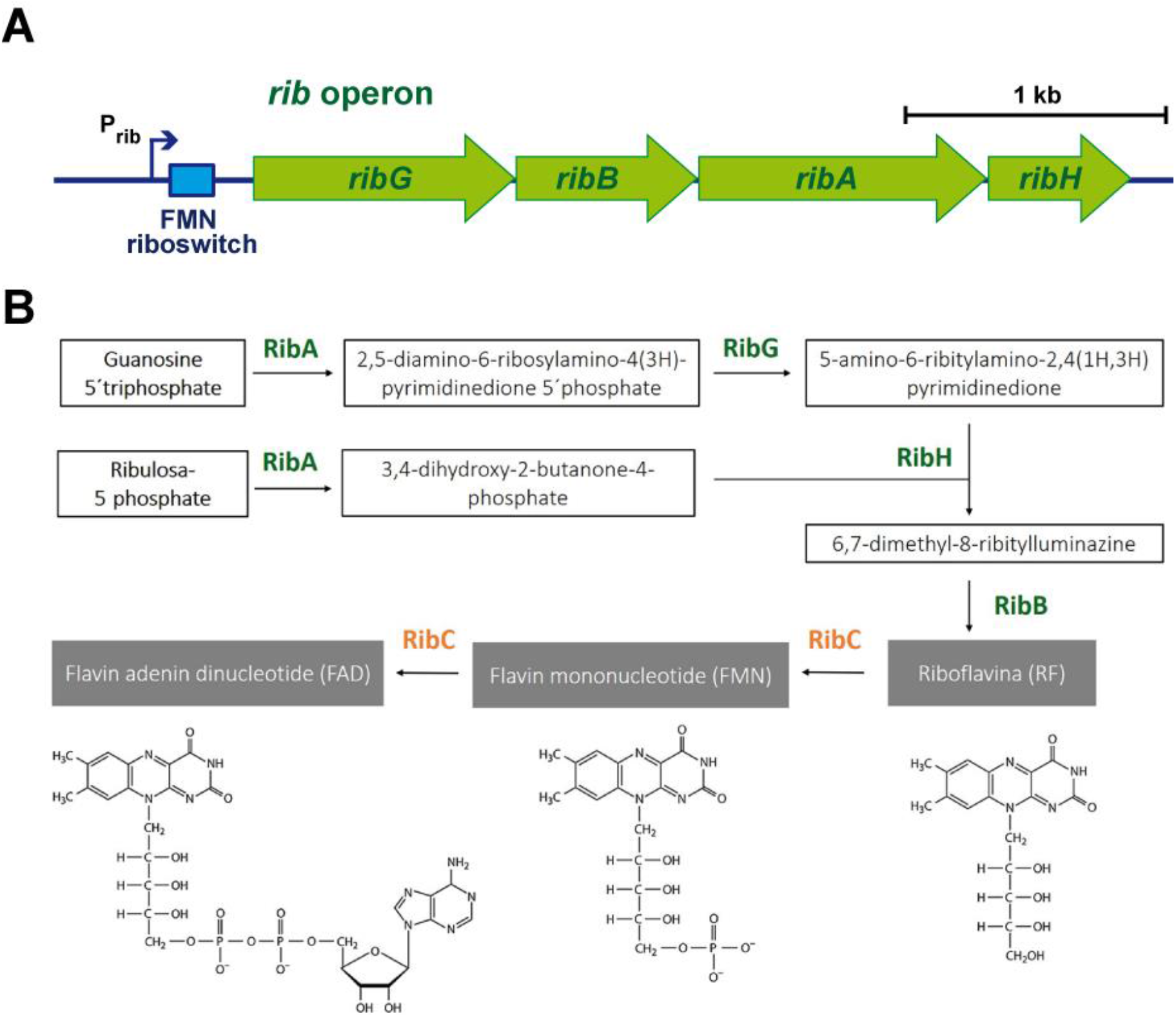
**A)** Expression of the *rib* operon for RF biosynthesis is initiated from the P_rib_ promoter and regulated by an FMN riboswitch (scheme adapted from (Capozzi *et al.*, 2012)). **B)** Overview of the riboflavin, FMN and FAD biosynthetic pathway. The enzymes encoded by the *rib* operon, which are involved in the RF biosynthesis, are indicated in green letters. RibG, riboflavin-specific deaminase and reductase; RibB, riboflavin synthase alpha subunit; RibA, a bifunctional enzyme which also catalyses the formation of 3,4-dihydroxy-2-butanone 4-phosphate from ribulose 5-phosphate; RibH, riboflavin synthase (beta subunit); RibC, bifunctional flavokinase/FAD synthetase, which is not encoded by the *rib* operon.

**Supporting Fig. S2.**
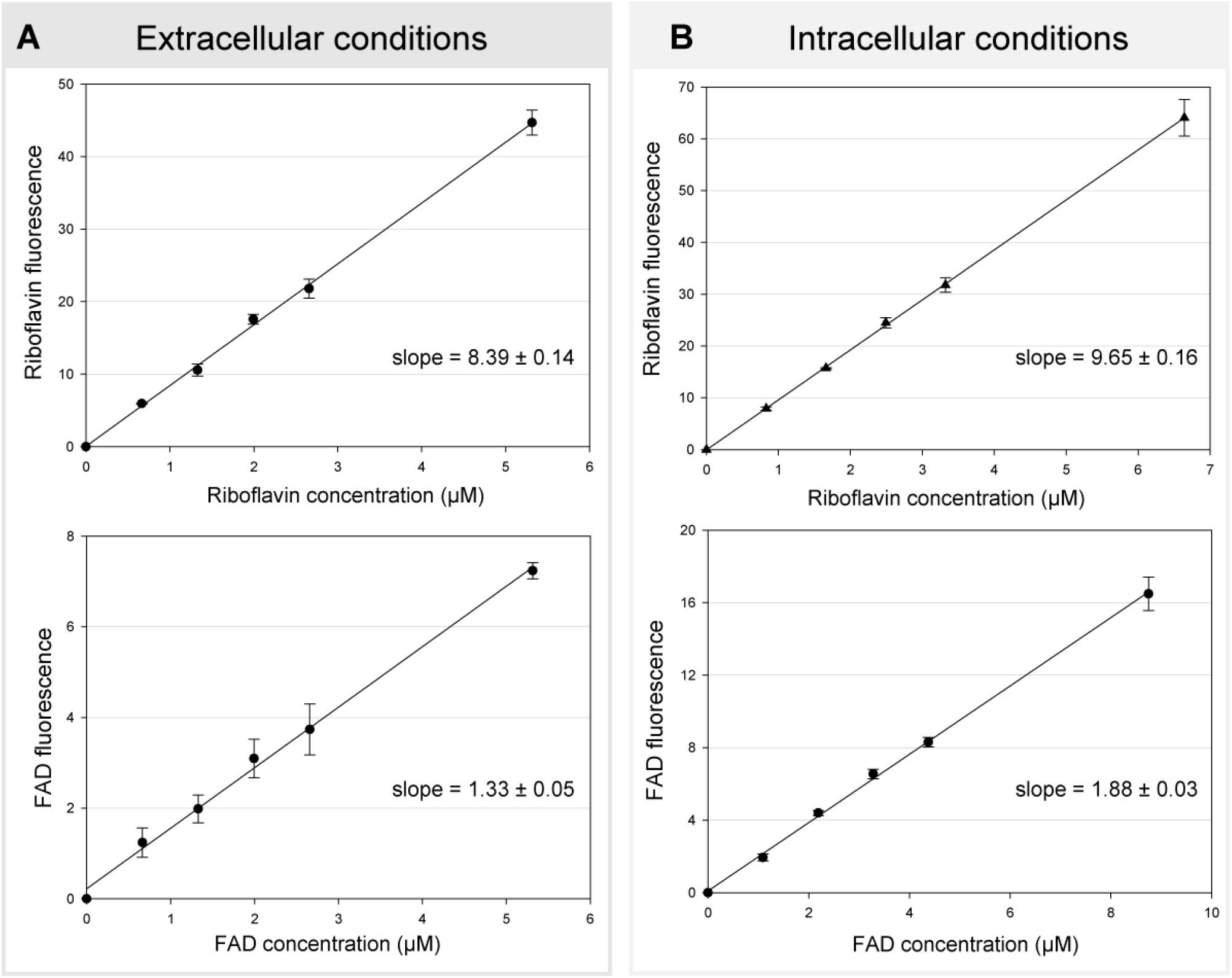
Calibration curves elaborated with RF and FAD solutions of defined concentrations prepared in media simulating the neutralized samples from the supernatant (**A**) and the pellet (**B**) of the cultures. The curves were obtained by using six different values of RF or FAD concentration and resulted in a highly linear relationship (R^2^ of 0.99). The experimental data were fitted to a linear regression model and the slope (intrinsic fluorescence of RF or FAD) is indicated in each panel. The calibration curves were elaborated from three independent experiments.

#### 13.2 Supporting Tables

**Supporting Table S1.**
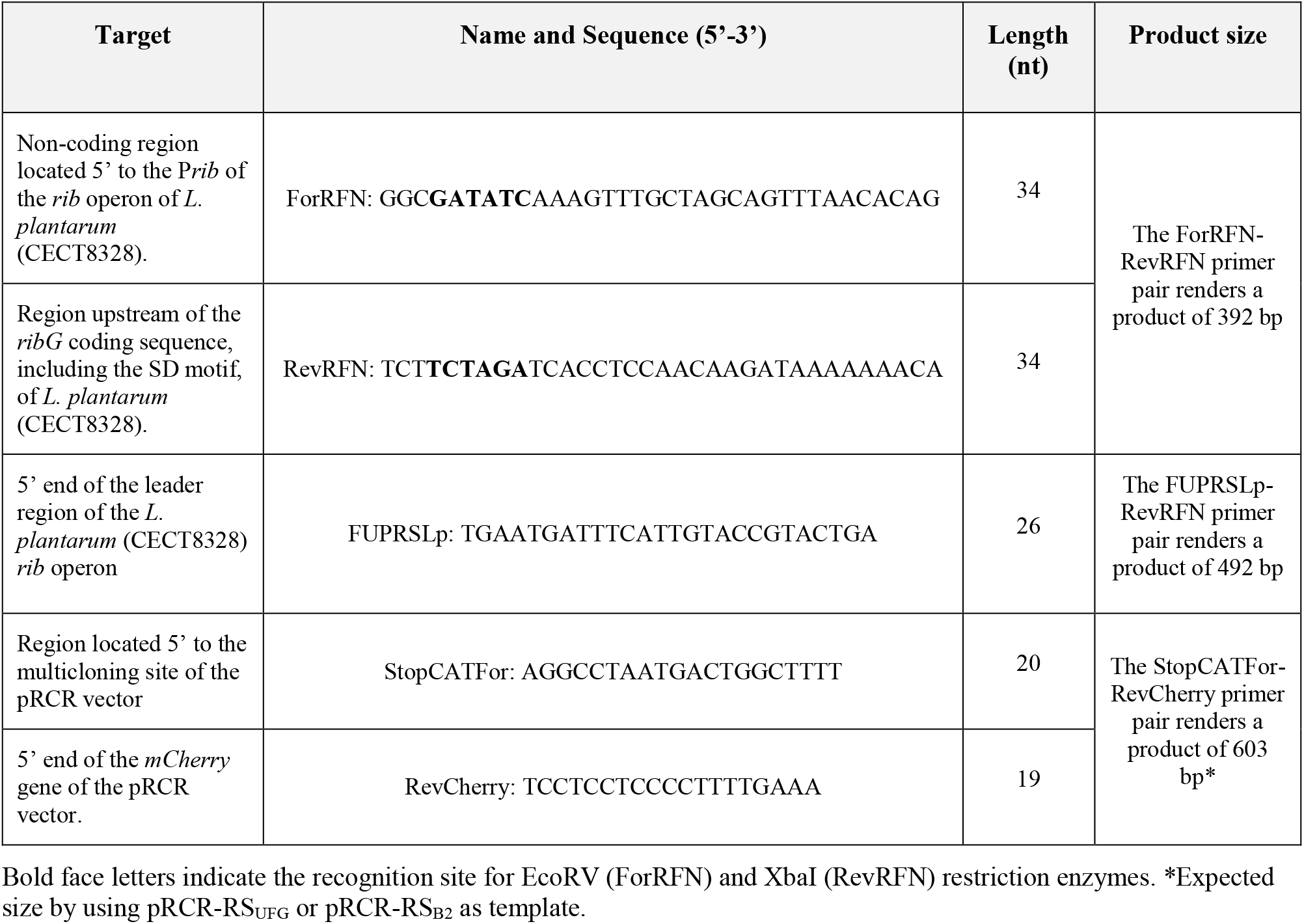
Oligonucleotides used in this study.

**Supporting Table S2.** Composition of the chemically defined medium (CDM) used in this work.

| Compound | Concentration (g/L) |
| --- | --- |
| CaCl<sub>2</sub> | $3.8 \times 10^{-2}$ |
| FeSO<sub>4</sub> | 0.02 |
| K<sub>2</sub>HPO<sub>4</sub> | 3.00 |
| KH<sub>2</sub>PO<sub>4</sub> | 3.00 |
| MgCl<sub>2</sub> | 0.20 |
| MnSO<sub>4</sub> | 0.01 |
| ZnSO<sub>4</sub> | $5 \times 10^{-3}$ |
| 4-aminobenzoic acid | $7.5 \times 10^{-3}$ |
| D-biotin | $3.8 \times 10^{-3}$ |
| Diammonium citrate | 0.60 |
| Sodium citrate anhydrous | 1.00 |
| Cyanocobalamin | $1.5 \times 10^{-3}$ |
| Folic acid | $1.5 \times 10^{-3}$ |
| Lipoic acid | $2.3 \times 10^{-3}$ |
| Nicotinic acid | $1.5 \times 10^{-3}$ |
| Ca-D-pantothenate | $1.5 \times 10^{-3}$ |
| Pyridoxine HCl | $1.5 \times 10^{-3}$ |
| Pyridoxamine 2 HCl | $3.8 \times 10^{-3}$ |
| Thiamine HCl | $1.5 \times 10^{-3}$ |
| Adenine sulphate | 0.01 |
| Guanine HCl | 0.01 |
| Xanthine | 0.01 |
| Thymine | 0.01 |
| Uracil | 0.01 |
| L-Alanine | 0.24 |
| L-Arginine | 0.12 |
| L-Asparagine | 0.35 |
| L-Aspartic acid | 0.41 |
| L-Cysteine-HCl | 0.27 |
| L-Glutamic acid | 0.49 |
| L-Glutamine | 0.38 |
| L-Glycine | 0.17 |
| L-Histidine | 0.15 |
| L-Leucine | 0.45 |
| L-Isoleucine | 0.20 |
| L-Lysine 2 HCl | 0.44 |
| L-Methionine | 0.12 |
| L-Phenylalanine | 0.27 |
| L-Proline | 0.67 |
| L-Serine | 0.34 |
| L-Threonine | 0.22 |
| L-Tryptophan | $4.8 \times 10^{-2}$ |
| L-Tyrosine | 0.29 |
| L-Valine | 0.32 |
| **Supplements** | **Concentration (%)** |
| Vitamin free casaminoacids | 0.10 |
| Glucose | 0.80 |

**Supporting Table S3.** Intra- or extracellular location of the real-time detected flavins.

| Sample | RF fluorescence |  |
| --- | --- | --- |
|  | *L. plantarum* UFG9 | *L. plantarum* B2 |
| Washed cells resuspended in fresh CDM | $0.90 \pm 0.12$ | $4.81 \pm 0.11$ |
| Supernatant | $0.94 \pm 0.03$ | $31.54 \pm 0.83$ |
| Washed cells resuspended in the culture supernatant | $2.52 \pm 0.14$ | $47.55 \pm 0.45$ |
| Uncentrifuged culture | $2.72 \pm 0.16$ | $49.27 \pm 0.70$ |

The values are expressed as mean ± experimental standard error of three measurements. The cellular pellets were washed twice with PBS 1X and resuspended in the same volume of either fresh CDM or their own collected supernatant. Triplicates (200-μL) of both cell suspensions, as well as of the supernatants and uncentrifuged cultures, were loaded in a 96-well optical bottom plate and analyzed for RF fluorescence.

## References

Abbas, C. A., and Sibirny, A. A. (2011). Genetic control of biosynthesis and transport of riboflavin and flavin nucleotides and construction of robust biotechnological producers. Microbiol. Mol. Biol. Rev. doi: 10.1128/mmbr.00030-10.

Acevedo-Rocha, C. G., Gronenberg, L. S., Mack, M., Commichau, F. M., and Genee, H. J. (2019). Microbial cell factories for the sustainable manufacturing of B vitamins. Curr. Opin. Biotechnol. doi:10.1016/j.copbio.2018.07.006.

Arena, M. P., Russo, P., Capozzi, V., López, P., Fiocco, D., and Spano, G. (2014). Probiotic abilities of riboflavin-overproducing *Lactobacillus* strains: A novel promising application of probiotics. Appl. Microbiol. Biotechnol. 98, 7569–7581. doi:10.1007/s00253-014-5837-x.

Berthier, F., Zagorec, M., Champomier-Vergès, M., Ehrlich, S. D., and Morel-Deville, F. (1996). Efficient transformation of *Lactobacillus sake* by electroporation. Microbiology 142, 1273–1279. doi:10.1099/13500872-142-5-1273.

Burgess, C. M., Smid, E. J., Rutten, G., and van Sinderen, D. (2006). A general method for selection of riboflavin-overproducing food grade micro-organisms. Microb. Cell Fact. 5, 24. doi:10.1186/1475-2859-5-24.

Burgess, C., O’Connell-Motherway, M., Sybesma, W., Hugenholtz, J., and Van Sinderen, D. (2004). Riboflavin production in *Lactococcus lactis:* Potential for in situ production of vitamin-enriched foods. Appl. Environ. Microbiol. 70, 5769–5777. doi:10.1128/AEM.70.10.5769-5777.2004.

Capozzi, V., Menga, V., Digesù, A. M., De Vita, P., Van Sinderen, D., Cattivelli, L., et al. (2011). Biotechnological production of vitamin B2-enriched bread and pasta. in Journal of Agricultural and Food Chemistry, 8013–8020. doi:10.1021/jf201519h.

Chen, J., Shen, J., Solem, C., and Jensen, P. R. (2013). Oxidative stress at high temperatures in *Lactococcus lactis* due to an insufficient supply of riboflavin. Appl. Environ. Microbiol. 79, 6140–6147. doi:10.1128/AEM.01953-13.

Darty, K., Denise, A., and Ponty, Y. (2009). VARNA: Interactive drawing and editing of the RNA secondary structure. Bioinformatics. doi:10.1093/bioinformatics/btp250.

Di Palma, F., Colizzi, F., and Bussi, G. (2013). Ligand-induced stabilization of the aptamer terminal helix in the *add* adenine riboswitch. RNA 19, 1517–1524. doi:10.1261/rna.040493.113.

Fabian, E., Bogner, M., Kickinger, A., Wagner, K. H., and Elmadfa, I. (2012). Vitamin status in elderly people in relation to the use of nutritional supplements. J. Nutr. Heal. Aging. doi: 10.1007/s12603-011-0159-5.

Garay-Novillo, J. N., García-Morena, D., Ruiz-Masó, J. Á., Barra, J. L., and del Solar, G. (2019). Combining modules for versatile and optimal labeling of lactic acid bacteria: two pMV158-family promiscuous replicons, a pneumococcal system for constitutive or inducible gene expression, and two fluorescent proteins. Front. Microbiol. 10. doi:10.3389/fmicb.2019.01431.

García-Angulo, V. A. (2017). Overlapping riboflavin supply pathways in bacteria. Crit. Rev. Microbiol. doi:10.1080/1040841X.2016.1192578.

Ge, Y. Y., Zhang, J. R., Corke, H., and Gan, R. Y. (2020). Screening and spontaneous mutation of pickle-derived *Lactobacillus plantarum* with overproduction of riboflavin, related mechanism, and food application. Foods 9, 88. doi:10.3390/foods9010088.

Juarez del Valle, M., Laiño, J. E., Savoy de Giori, G., and LeBlanc, J. G. (2014). Riboflavin producing lactic acid bacteria as a biotechnological strategy to obtain bio-enriched soymilk. Food Res. Int. 62, 1015–1019. doi:10.1016/j.foodres.2014.05.029.

Kang, C., Wu, H. L., Xu, M. L., Yan, X. F., Liu, Y. J., and Yu, R. Q. (2019). Simultaneously quantifying intracellular FAD and FMN using a novel strategy of intrinsic fluorescence four-way calibration. Talanta 197, 105–112. doi:10.1016/j.talanta.2018.12.076.

Kimata, S., Mochizuki, D., Satoh, J., Kitano, K., Kanesaki, Y., Takeda, K., et al. (2018). Intracellular free flavin and its associated enzymes participate in oxygen and iron metabolism in *Amphibacillus xylanus* lacking a respiratory chain. FEBS Open Bio 8, 947–961. doi:10.1002/2211-5463.12425.

Lacks, S. A. (1968). Genetic regulation of maltosaccharide utilization in pneumococcus. Genetics 60, 685–706.

Lacks, S. A., and Greenberg, B. (1977). Complementary specificity of restriction endonucleases of *Diplococcus pneumoniae* with respect to DNA methylation. J. Mol. Biol. 114, 153–168.

LeBlanc, J. G., Milani, C., de Giori, G. S., Sesma, F., van Sinderen, D., and Ventura, M. (2013). Bacteria as vitamin suppliers to their host: a gut microbiota perspective. Curr. Opin. Biotechnol. 24, 160–168. doi:10.1016/j.copbio.2012.08.005.

López, P., Espinosa, M., Stassi, D. L., and Lacks, S. A. (1982). Facilitation of plasmid transfer in *Streptococcus pneumoniae* by chromosomal homology. J. Bacteriol. 150, 692–701.

Magnúsdóttir, S., Ravcheev, D., De Crécy-Lagard, V., and Thiele, I. (2015). Systematic genome assessment of B-vitamin biosynthesis suggests cooperation among gut microbes. Front. Genet. doi:10.3389/fgene.2015.00148.

Massey, V. (2000). The chemical and biological versatility of riboflavin. in Biochemical Society Transactions doi:10.1042/bst0280283.

Mazur-Bialy, A. I., Pochec, E., and Plytycz, B. (2015). Immunomodulatory effect of riboflavin deficiency and enrichment – Reversible pathological response versus silencing of inflammatory activation. J. Physiol. Pharmacol.

McAnulty, M. J., and Wood, T. K. (2014). YeeO from *Escherichia coli* exports flavins. Bioengineered 5, 386–392. doi: 10.4161/21655979.2014.969173.

Mensink, G. B. M., Fletcher, R., Gurinovic, M., Huybrechts, I., Lafay, L., Serra-Majem, L., et al. (2013). Mapping low intake of micronutrients across Europe. Br. J. Nutr. doi:10.1017/S000711451200565X.

Mohedano, M. L., García-Cayuela, T., Pérez-Ramos, A., Gaiser, R. A., Requena, T., and López, P. (2014). Construction and validation of a mCherry protein vector for promoter analysis in *Lactobacillus acidophilus*. J. Ind. Microbiol. Biotechnol. doi:10.1007/s10295-014-1567-4.

Mohedano, M. L., Hernández-Recio, S., Yépez, A., Requena, T., Martínez-Cuesta, M. C., Peláez, C., et al. (2019). Real-time detection of riboflavin production by *Lactobacillus plantarum* strains and tracking of their gastrointestinal survival and functionality *in vitro* and *in vivo* using mCherry labeling. Front. Microbiol. 10, 1–13. doi:10.3389/fmicb.2019.01748.

Pinto, J., Huang, Y. P., and Rivlin, R. S. (1987). Mechanisms underlying the differential effects of ethanol on the bioavailability of riboflavin and flavin adenine dinucleotide. J. Clin. Invest. 79, 1343–1348. doi:10.1172/JCI112960.

Revuelta, J. L., Ledesma-Amaro, R., Lozano-Martinez, P., Díaz-Fernández, D., Buey, R. M., and Jiménez, A. (2017). Bioproduction of riboflavin: a bright yellow history. J. Ind. Microbiol. Biotechnol. 44, 659–665. doi:10.1007/s10295-016-1842-7.

Ruiz-Cruz, S., Solano-Collado, V., Espinosa, M., and Bravo, A. (2010). Novel plasmid-based genetic tools for the study of promoters and terminators in *Streptococcus pneumoniae* and *Enterococcus faecalis*. J. Microbiol. Methods 83, 156–163. doi:10.1016/j.mimet.2010.08.004.

Russo, P., Capozzi, V., Arena, M. P., Spadaccino, G., Dueñas, M. T., López, P., et al. (2014a). Riboflavin-overproducing strains of *Lactobacillus fermentum* for riboflavin-enriched bread. Appl. Microbiol. Biotechnol. 98, 3691–3700. doi:10.1007/s00253-013-5484-7.

Russo, P., de Chiara, M. L. V., Capozzi, V., Arena, M. P., Amodio, M. L., Rascón, A., et al. (2016). *Lactobacillus plantarum* strains for multifunctional oat-based foods. LWT – Food Sci. Technol. doi:10.1016/j.lwt.2015.12.040.

Russo, P., De Chiara, M. L. V., Vernile, A., Amodio, M. L., Arena, M. P., Capozzi, V., et al. (2014b). Fresh-cut pineapple as a new carrier of probiotic lactic acid bacteria. Biomed Res. Int. doi:10.1155/2014/309183.

Russo, P., Iturria, I., Mohedano, M. L., Caggianiello, G., Rainieri, S., Fiocco, D., et al. (2015a). Zebrafish gut colonization by mCherry-labelled lactic acid bacteria. Appl. Microbiol. Biotechnol. doi:10.1007/s00253-014-6351-x.

Russo, P., Peña, N., de Chiara, M. L. V., Amodio, M. L., Colelli, G., and Spano, G. (2015b). Probiotic lactic acid bacteria for the production of multifunctional fresh-cut cantaloupe. Food Res. Int. 77, 762–772. doi:10.1016/j.foodres.2015.08.033.

Serganov, A., Huang, L., and Patel, D. J. (2009). Coenzyme recognition and gene regulation by a flavin mononucleotide riboswitch. Nature 458, 233–237. doi:10.1038/nature07642.

Sherwood, R. A. (2018). “Methods for assessment of vitamin B2,” in Laboratory Assessment of Vitamin Status (Elsevier), 165–172. doi:10.1016/B978-0-12-813050-6.00007-3.

Thakur, K., Tomar, S. K., and De, S. (2016). Lactic acid bacteria as a cell factory for riboflavin production. Microb. Biotechnol. 9, 441–451. doi:10.1111/1751-7915.12335.

Wickiser, J. K., Winkler, W. C., Breaker, R. R., and Crothers, D. M. (2005). The speed of RNA transcription and metabolite binding kinetics operate an FMN riboswitch. Mol. Cell. doi: 10.1016/j.molcel.2005.02.032.

Winkler, W. C., Cohen-Chalamish, S., and Breaker, R. R. (2002). An mRNA structure that controls gene expression by binding FMN. Proc. Natl. Acad. Sci. 99, 15908–15913. doi:10.1073/pnas.212628899.

Yépez, A., Russo, P., Spano, G., Khomenko, I., Biasioli, F., Capozzi, V., et al. (2019). *In situ* riboflavin fortification of different kefir-like cereal-based beverages using selected Andean LAB strains. Food Microbiol. doi: 10.1016/j.fm.2018.08.008.

Zheng, J., Wittouck, S., Salvetti, E., Franz, C. M. A. P., Harris, H. M. B., Mattarelli, P., et al. (2020). A taxonomic note on the genus *Lactobacillus*: Description of 23 novel genera, emended description of the genus *Lactobacillus* Beijerinck 1901, and union of *Lactobacillaceae* and *Leuconostocaceae*. Int. J. Syst. Evol. Microbiol. 70, 2782–2858. doi:10.1099/ijsem.0.004107.

Zhu, Y. Y., Thakur, K., Feng, J. Y., Cai, J. S., Zhang, J. G., Hu, F., et al. (2020). Riboflavin-overproducing lactobacilli for the enrichment of fermented soymilk: insights into improved nutritional and functional attributes. Appl. Microbiol. Biotechnol. doi:10.1007/s00253-020-10649-1.

## Supporting References

Capozzi, V., Russo, P., Dueñas, M.T., López, P., and Spano, G. (2012) Lactic acid bacteria producing B-group vitamins: A great potential for functional cereals products. Appl Microbiol Biotechnol.

